# Mechanical state of pre-tumoral epithelia dictates subsequent tumour aggressiveness

**DOI:** 10.1101/2024.05.17.594669

**Authors:** Marianne Montemurro, Bruno Monier, Magali Suzanne

## Abstract

Tumours evolve through the acquisition of increasingly aggressive traits associated with dysplasia. This progression is accompanied by the alteration of tumour mechanical properties, especially through extracellular matrix remodelling. However, the contribution of intrinsic tumour cells mechanics to tumour aggressiveness remains unknown *in vivo*.

Here we show that adherens junction tension in pre-tumoral tissues dictates subsequent tumour evolution. Increased cell contractility, observed in aggressive tumours before any sign of tissue overgrowth, proved sufficient to trigger dysplasia in normally hyperplastic tumours. Unexpectedly, high contractility contributes to tumour evolution through cell death induction which favours cell-cell junctions weakening. Overall, our results highlight the need to re-evaluate the roles of tumoral cell death, and identify pre-tumour cell mechanics as an unsuspected early marker and key trigger of tumour aggressiveness.

**GRAPHICAL ABSTRACT:** 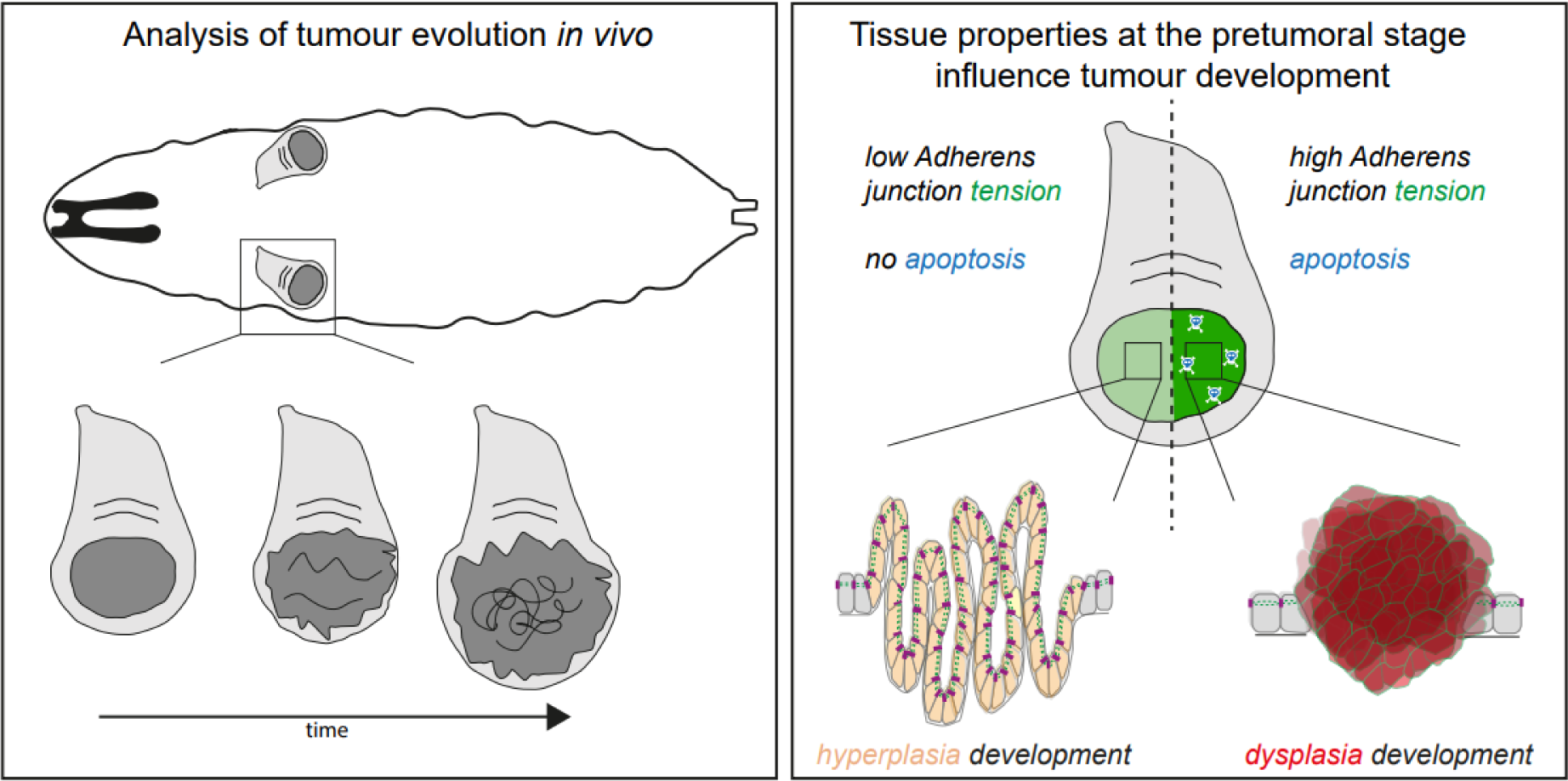

## INTRODUCTION

Cancer is a highly complex pathology leading to tissue overgrowth and, in most dramatic cases, to colonization of distant organs. Cancer is characterized by several hallmarks, such as sustained proliferative properties, evasion from immunological monitoring or increased resistance to cell death^1^. A vast majority of tumours, carcinoma, stems from epithelial cells. Epithelia are sheets of polarized cells, presenting an apico-basal polarity, connected among each other through intercellular junctions, especially adherens junctions (that sustain mechanical integrity of the tissue) and tight junctions (that control permeability). Epithelial cells are also connected via receptors such as Integrin to the extracellular matrix (ECM) that provides structural stability ^2^. Tumours can evolve through increasingly aggressive steps ^3–5^, in part due to the acquisition of mesenchymal traits such as loss of apico-basal polarity and decrease in intercellular or ECM adhesion. Thus, tumour aggressiveness is associated with an epithelial-to-mesenchymal transition (EMT), a process which can be partial, leading to cells expressing a combination of epithelial and mesenchymal markers ^6,7^. Tumoral EMT therefore underlies the switch from initially epithelial, hyperplastic to more aggressive dysplastic tumours ^8^. Because appearance of mesenchymal domains also favours the formation of cancer stem cells and increases resistance to therapeutic strategies ^9–11^, understanding mechanisms initiating cancer-associated EMT is therefore of prime importance.

One striking characteristic of tumours of epithelial origins is the profound alteration in their mechanical properties. For instance, tissue stiffness is dramatically increased in tumours, and several mechanisms underly this change in properties, such as constrained growth ^12^, increase in interstitial fluid pressure ^13^, or increase in ECM deposition or crosslinking ^14,15^. Despite acknowledgement of the importance of biomechanics associated with tumour evolution, the contribution of tumour cell mechanics is still unclear, especially because some inconsistency exists in the data obtained so far on tumour cell physical properties. On the one hand, the vast majority of rheological studies analysing the stiffness of tumour cells in culture described them as softer than normal cells, leading to the hypothesis of an inverse correlation between cell stiffness and malignancy ^16,17^. On the other hand, an increase in cellular contractility has been associated with tumour growth in breast cancer cells, mouse skin or *Drosophila* eye epithelia ^18–20^. These studies suggest that cancer cell stiffness could contribute to tumour development, although mechanical properties were not directly analysed in those contexts. The discrepancy between data probing mechanical properties of tumour cells may be the consequence of i-the difficulty to discriminate between tumour cells and their microenvironment in *in situ* studies and ii-the non-physiological conditions associated with tumour cells in culture. Overall, these controversial data highlight the need to characterize the evolution of tumour mechanics throughout tumour progression in a simplified and integrated *in vivo* system. For this purpose, we performed a detailed characterization of actomyosin cytoskeleton dynamics and contribution in temporally and spatially controlled *Drosophila* tumours, using an inducible system to follow tumour development.

## RESULTS

### Highly stereotyped development of dysplastic tumours in the *Drosophila* wing disc

To probe tissue mechanics in evolving tumours, we used the downregulation of the tumour-suppressor gene *Syntaxin 7* (*Syx7*, also known as *Avalanche*, *Avl*), a component of the endocytic machinery that controls trafficking of transmembrane molecules such as the Notch receptor and which is associated with dysplasia in the fly follicular epithelium ^21^. We generated tumours in the pouch region of the *Drosophila* wing disc since this domain, in which transgene expression can be timely controlled, is composed of a simple monolayer epithelium sitting on the extracellular matrix. This system allows to unambiguously probe the mechanical properties of tumour cells, without any possible confusion with cells of the microenvironment. We confirmed that Syx7 knockdown can promote tumour formation in the wing pouch ^22,23^ (Sup Fig.1A) and performed a detailed characterization of tumour evolution at cell and tissue scales in this context.

**Figure 1.**
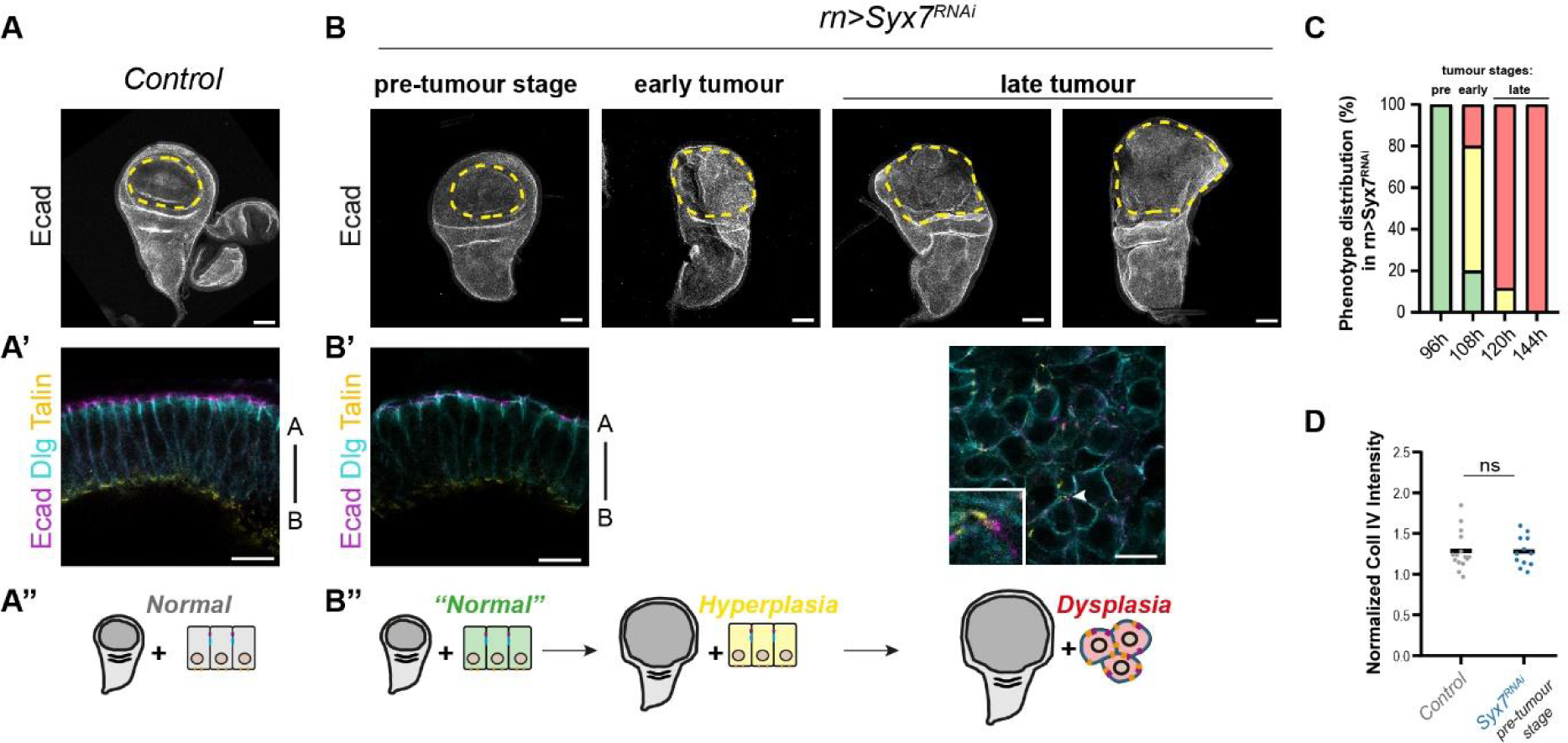
Syx7 RNAi tumour evolution is progressive and stereotyped in the wing disc. (A, B) E-Cad staining of control (A) and Syx7 RNAi (B) wing discs (projections), showing overall tissue morphology. The region of Syx7 RNAi expression, the wing pouch, is delineated by a dotted yellow line. In (B), discs shown from left to right were collected at 96h, 108h, 120h and 144h after egg laying (AEL). Note the progressive increase in size of the pouch domain in Syx7 RNAi discs from 108h onwards. The control disc shown in (A) was collected at 96h AEL. (A’, B’) Single plane close up showing organisation of wing pouch tissues at the 96h AEL from control (A’) and Syx7 RNAi disc (B’, left), and at 120h after tumour formation (B’, right). Note that cells were epithelial at 96h, but have become mesenchymal at 120h AEL in Syx7 RNAi tumours, with apical (ECad and Dlg, magenta and cyan) and basal (Talin, yellow) structures in close vicinity (see inset taken at the level of the arrowhead), and cells becoming round. (A’’, B’’) Schemes summarizing global tissue size and tissue organization (epithelial vs mesenchymal) as illustrated in (A-B’). Stages (normal, hyperplasia, dysplasia) are therefore determined based on the size of the wing pouch and the cellular organisation. (C) Histogram showing the distribution of wing disc phenotypes at the indicated time points (+/-1.5h), in Syx7 RNAi tumours according to the legend in (B’’). Number of discs analysed at 96, 108, 120h and 144h AEL are respectively 46, 15, 26, 31. (D) Analysis of extracellular matrix intensity in control (grey, n=15) and Syx7 RNAi (blue, n=12) at the pre-tumour stage (96h AEL). The intensity of Collagen IV was quantified on sagittal sections (as shown in Sup Fig.1D). On the graph, each dot corresponds to the level of Collagen IV expression in one individual disc. In all panels, transgenes are driven in the wing pouch using the *rotund*::Gal4 driver at 25°C. Scale bars: 50µm for panels A, B and 10µm for panels A’, B’. Apical-basal (A-B) is indicated for each close up showing a polarized epithelial tissue. Statistical tests: Mann-Whitney for D. p-value: ns, non-significant.

We combined the macroscopic description of tissue growth (Fig. 1A-B) with detailed characterization of tissue organization using a combination of markers for adherens junctions (E-cadherin, E-Cad), tight junctions (Disc large, Dlg), cell/ECM junctions (Talin, Beta-Integrin) (Fig.1A’-B’). We found that Syx7 RNAi discs were initially similar or slightly smaller than control discs at 96h after egg laying (AEL), with E-Cad/Dlg and Talin respectively labelling apical and basal poles of epithelial cells (Fig.1B’). Since no sign of tumour development is visible at this stage, we call it “pre-tumoral stage”. At 108h AEL, a stage we call “early tumour stage”, most Syx7 RNAi discs are overgrown compared to control discs of the same age, still keeping epithelial organization, corresponding to hyperplastic tumours. Finally, by 120h AEL onwards, which we call the “late tumour stage”, nearly all Syx7 RNAi discs have become dysplastic. Indeed, these tumoral tissues are not only overgrown but also present domains in which cells have lost their epithelial organization, as revealed by a loose shape, Dlg relocation around most of the cell cortex, and scattered E-Cad and Talin dots (Fig.1B’, right panel). Similar results regarding the loss of epithelial polarity in late tumours were observed using additional markers, such as the basal beta-Integrin, the apical cell polarity marker Bazooka/Par3 and F-actin which labels cortex organisation (Sup Fig. 1B, C), revealing a mesenchymal-like state (hereafter referred to as “mesenchymal”). Hence, at 25°C, we identified three main stages, with the “pre-tumour stage” at 96h AEL, just before the onset of overgrowth, the “early tumour stage” at 108h AEL in which most discs are hyperplastic, and the late tumour stage from 120h AEL onwards in which nearly all tumours are dysplastic (Fig.1 B’’, C).

### Increased adherens junction contractility is a marker of pre-tumoral tissues in evolving tumours

This temporal characterization of tumour evolution set up the framework to investigate possible alterations of tissue mechanics in evolving tumours. We started studying tumour development starting from the pre-tumoral stage. Since part of tissue mechanics relies on ECM constitution, we first characterized the pattern of one main ECM component, Collagen IV, as well as ECM/tumour cell interaction, by looking at the mechanoresponsive component of Integrin junctions, Talin. Neither Collagen IV levels nor Talin distribution were different between control and Syx7 RNAi discs at the pre-tumour stage (compare Fig. 1A’ with the left panel in Fig1B’, Fig. 1D, Sup Fig. 1D). Next, we focused on the cytoskeleton itself, which could be responsible for intrinsic alterations of tumour cell mechanics. Strikingly, we observed an unsuspected strong enrichment of F-actin and of the molecular motor non-muscle myosin II (hereafter referred to as myosin II) in Syx7 RNAi discs at the pre-tumour stage (Fig. 2A and Sup Fig.1E). This pointed towards a potential alteration of tumour cell mechanics already in pre-tumoral tissues. We quantified the levels of F-actin and myosin II, both on the basal side where basal junctions connect cells to the extracellular matrix, and on the apical side at the level of intercellular junctions. Interestingly, both F-actin and myosin II were strongly enriched apically (Fig. 2C, D). Indeed, while F-actin levels were not significantly different at the basal side and myosin II levels were increased by 15%, F-actin and myosin II were increased by about respectively 25% and nearly 50% at the apical pole of pre-tumoral tissue. Co-staining with E-cadherin showed that F-actin and myosin II accumulate specifically at adherens junctions (Fig. 2B), suggesting an unexpected change in adherens junction contractility between a control and a pre-tumoral tissue.

**Figure 2.**
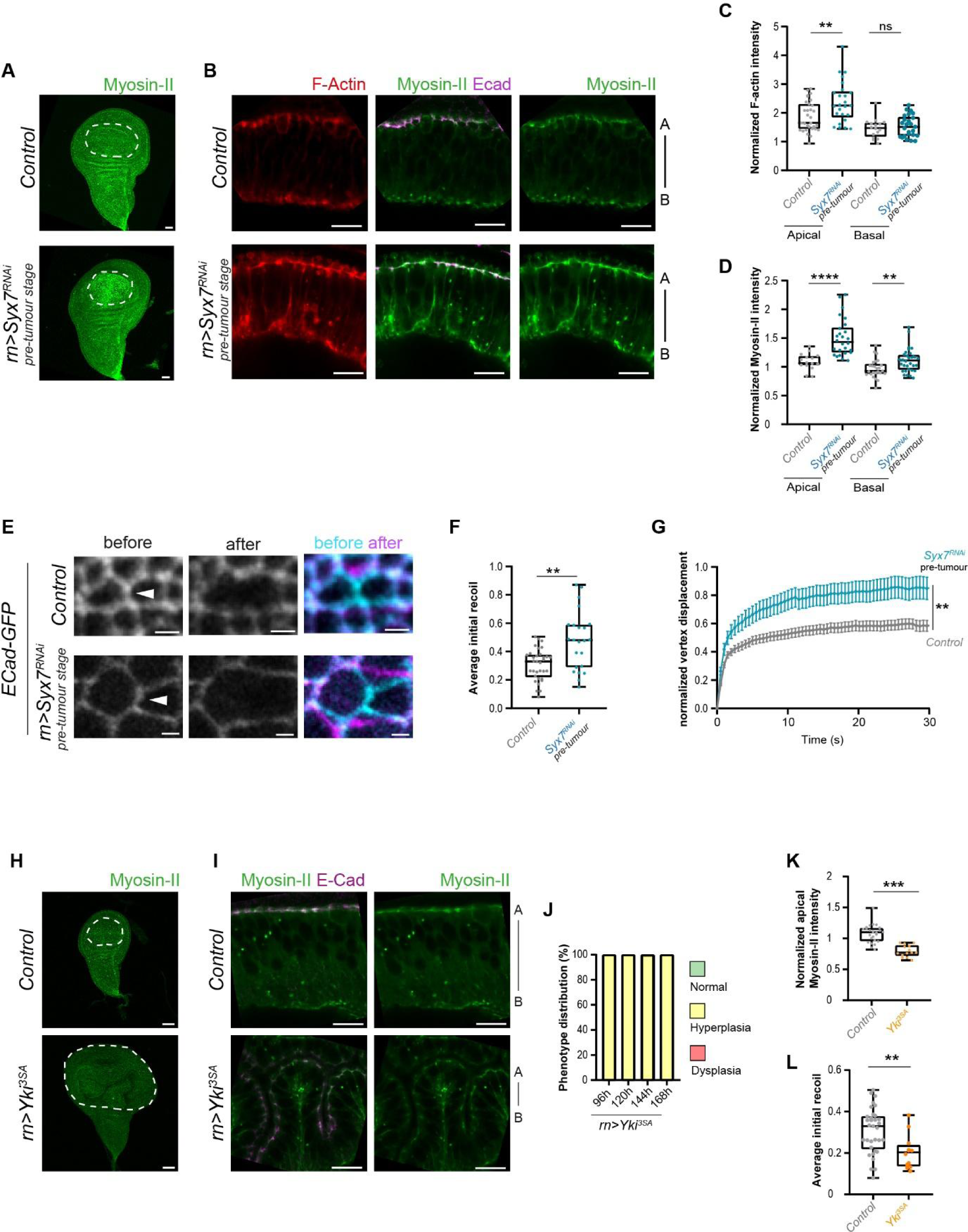
Increased adherens junction contractility constitutes an early marker of aggressive tumours. (A) Wing discs expressing endogenous myosin II-GFP (green) at the pre-tumour stage (96h AEL), showing myosin II accumulation in the pouch (white dashed line) following Syx7 RNAi knockdown (bottom) compared to the control (top). (B) Close up views showing F-actin (red), myosin II (green) and E-cadherin (magenta) in control (top) and Syx7 RNAi (bottom) epithelial tissues at the pre-tumour stage (96h AEL). Note the accumulation of F-actin and myosin II at adherens junctions before any sign of tumour in Syx7 RNAi. (C, D) Box plots showing F-actin (C) and myosin II (D) fluorescence levels at adherens junctions (apical) and basal junctions (basal) in the wing pouch of control (grey) and Syx7 RNAi (blue) discs at the pre-tumour stage (96h AEL). Pouch levels are normalized with levels in the unmanipulated hinge/notum region for each disc. Numbers of discs analysed in control and Syx7 RNAi discs are respectively 31 and 25 in apical and 16 and 25 in basal for (C) and 40 and 32 in apical and 25 and 32 in basal for (D). (E-G) Analysis of adherens junction tension by laser ablation in control and Syx7 RNAi discs expressing E-Cadherin-GFP at the pre-tumour stage (96h AEL). (E) Example of single plane images before and after laser ablation. (F) Box plots showing quantification of the initial recoil of junctions from control (grey, n=33) and Syx7 RNAi (blue, n=23). (G) Curves of normalized vertex displacement following laser ablation in the same genetic backgrounds. Number of junctions analysed: 33 and 29 in control and Syx7 RNAi respectively. (H) Wing discs expressing endogenous myosin II-GFP (green) at 96h AEL, in control (top) and Yorkie overexpression (bottom). Myosin II does not accumulate in the pouch (white dashed line) following Yorkie overexpression. (I) Close up views showing myosin II (green) and E-cadherin (magenta) in control (top) and Yorkie overexpression (bottom) contexts (96h AEL). Note epithelial convolutions associated with tissue hyperplasia in Yorkie tumours. (J) Histogram showing the distribution of wing disc stages at the indicated time points (+/-1.5h) in Yorkie tumours. Number of discs analysed at 96h, 120h, 144h, 168h AEL are respectively 47, 43, 18, 12. (K) Box plots showing myosin II fluorescence levels at adherens junctions in the wing pouch of control (grey, n= 24) and Yorkie overexpression (orange, n=14) contexts at 96h AEL. (L) Analysis of adherens junction tension by laser ablation in control (grey) and Yorkie overexpression (orange) contexts expressing E-Cadherin-GFP at 96h AEL. Box plots shows quantification of the initial recoil of junctions (n=33 in control and 12 in Yorkie). Same controls are shown in (F) and in (L). In all panels, transgenes are driven in the wing pouch using the *rotund*::Gal4 driver at 25°C. Scale bars: 10µm, except 50µm for panels A, H and 2µm for panel E. Apical-basal (A-B) is indicated for each close up showing a polarized epithelial tissue. In box plots, the black bar indicates the median, the whiskers indicate the maximal range, and each value is indicated by a dot. Statistical tests: T-test for (C, F, G, K) and Mann-Whitney for D, L). p-value: **, <0.01; ***, <0.001; ****, <0.0001, ns, non-significant.

We therefore determined whether acto-myosin accumulation efficiently translates into an alteration of adherens junction’s tension. For this purpose, we performed laser ablation on the E-cadherin-GFP knock-in line in control and Syx7 RNAi discs at pre-tumour stage (Fig. 2E). Interestingly, the recoil of vertices following ablation is higher in the Syx7 RNAi discs, with both initial and maximal recoils being about 50% higher compared to control discs (Fig. 2F, G). The increase of adherens junction contractility is observed at the pre-tumour stage, when the tumour is not yet morphologically identifiable as an overgrowth. It thus appears to be the first sign of tumour development in this context and constitutes a pre-tumoral marker.

We next wondered whether the increase in adherens junction tension is specific to aggressive tumours, or whether it is also observed in early hyperplastic tumours. We therefore turned to a genetic context strictly associated with hyperplasia induction in fly: the overexpression of Yorkie, the ortholog of the mammalian YAP transcription factor ^24^. Following expression of a mutated form of Yorkie ^25^ in the wing discs, we observed that at any timepoint analysed the discs are larger than the control ones (Fig. 2H, Sup Fig.2A, B) and that cells keep an epithelial organization (Fig.2I, Sup Fig.2C, D). Therefore, EMT is never induced following Yorkie expression and tumours remain hyperplastic, irrespective of their age (Fig. 2J). Importantly, myosin II levels were not increased at adherens junctions in this context (Fig.2I, K), and their tension was even slightly decreased (Fig.2L, Sup Fig. 2E). Thus, comparing analysis in dysplastic (above) and hyperplasic contexts, we conclude that high adherens junction tension constitutes a pre-tumoral marker specific of dysplastic tumours.

### High junctional contractility in tumour cells drives tumour aggressiveness

We then asked whether high tension could promote tumour evolution and aggressiveness. We therefore tested whether increasing junctional contractility could be sufficient to promote an epithelial-to-mesenchymal cell behaviour in strictly hyperplastic tumours, such as Yorkie tumours.

To increase contractility at adherens junction, we set up two independent strategies, initially in non-tumoral wing discs, to ensure that these genetic manipulations do not perturb epithelial organisation. First, we induced the specific recruitment of myosin II at adherens junctions through the use of a trap forcing GFP-fusion protein relocation. This tool, hereafter called AJ-GFP-trap (Fig.3A) is composed of a llama antibody (or nanobody) against GFP fused to the adherens junction protein alpha-Catenin that controls its location and to a red fluorescent protein (RFP) to visualize it ^26,27^. When the myosin II regulatory light chain (MRLC) *spaghetti squash* (*sqh*) endogenously tagged by GFP ^28^ is expressed in the wing pouch, the AJ-GFP-trap leads to a strong increase of myosin II at the apex of epithelial cells from the wing pouch (Fig.3B, C, Sup Fig.3A, B). Analysis of myosin II intensity shows that it increases by 50% at adherens junctions in presence of the AJ-GFP-trap by comparison with a control transgene expressing simply an alpha-catenin-RFP construct (Fig.3C), reaching levels close to those observed in Syx7 RNAi discs (Fig.2D). In a complementary way to increase contractility, we inactivated the *flapwing* gene product, a component of the phosphatase complex that restrain myosin II activity through dephosphorylation of *spaghetti squash* (Sup Fig.3C) ^29,30^. *flapwing* knockdown also led to an increase of myosin II in the wing pouch by 20% at adherens junction’s, but no significantly increase at the basal pole (Sup Fig.3D-F). The increase in apical myosin II levels translated in enhanced vertex recoil following laser ablation, revealing enhanced adherens junction’s tension in this context (Fig.3D, Sup Fig. 3G). Thus, these strategies induce higher myosin II recruitment and increased tension at adherens junctions in the wing disc without affecting epithelial organization in non-tumoral epithelium, as revealed by Dlg, alpha-catenin and myosin II normal distribution (Fig.3B, Sup Fig.3B, E).

**Figure 3.**
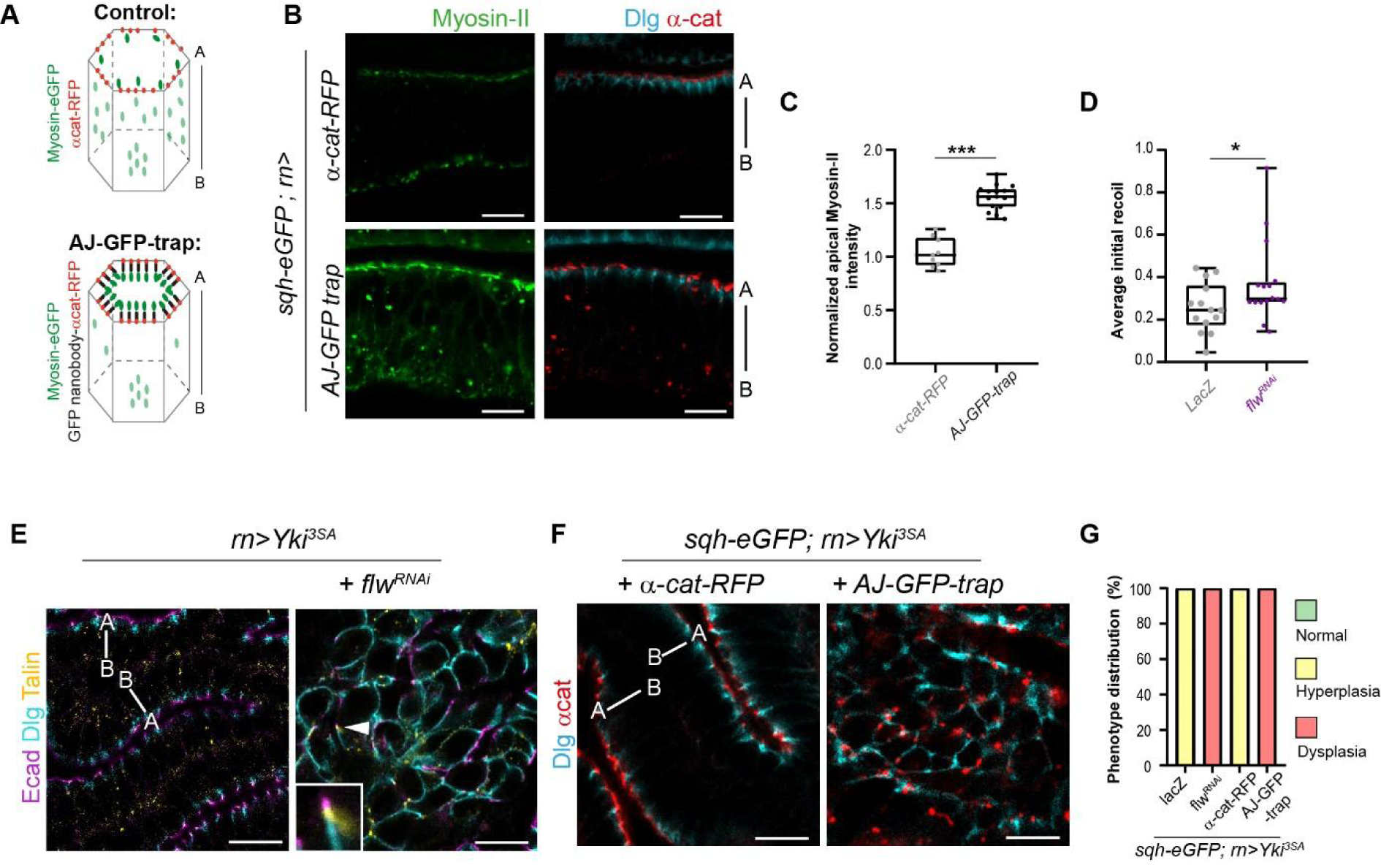
High tumour cell contractility triggers dysplasia. (A) Principle of AJ-GFP-trap function to target myosin II-GFP accumulation specifically at adherens junctions. Cytoplasmic and adherens junctions’ pools of myosin II are respectively shown in light and dark green. (B-D) Analysis of wing discs mechanics in late third instar larvae. (B) Close up views of the epithelial tissue showing myosin II (green) and adherens junctions (red). Note the strong enrichment of myosin II at adherens junctions caused by the presence of the AJ-GFP trap (bottom), compared to the control (top). Dlg (cyan) localisation is still localised apically following enrichment of myosin II caused by the trap (bottom). (C) Quantification of myosin II-GFP levels at adherens junctions of the wing pouch in control discs (expressing alpha-catenin-RFP, n=9), or following expression of the AJ-GFP trap fused to alpha-catenin-RFP (n=15). (D) Analysis of adherens junction tension by laser ablation in control (expressing lacZ, grey) and flapwing RNAi discs (magenta) co-expressing E-Cadherin-GFP. Box plots show quantification of the initial recoil of junctions following laser ablation. Number of junctions analysed: 14 and 16 in control and flapwing RNAi respectively. (E) Close up views of the organisation of Yorkie tumours without (control, left, n=27) or with myosin II increase at adherens junction due to flapwing RNAi knockdown (right, n=4). Cell rounding and loss of epithelial organisation is associated with loss of segregation of apico-basal polarity makers, as shown by E-Cadherin/Dlg/Talin proximity in the inset taken at the level of the arrowhead. (F) Close up views of the organisation of Yorkie tumours without (alpha-Catenin-RFP expression, left, n=30) or with myosin II increase at adherens junction due to expression of the AJ-GFP trap (right, n=17). (G) Histogram showing the distribution of Yorkie tumours co-expressing the indicated transgenes. Numbers of discs analysed: alpha-cat-RFP (control), 30; AJ-GFP-Trap, 17; lacZ (control), 14; *flapwing* RNAi, 16. Transgenes are driven in the wing pouch using the *rotund*::Gal4 driver at 25°C, apart from panels (D) where pdm2::Gal4 was used. Experiments with the AJ-GFP trap were conducted at 22°C while experiments with *flapwing* RNAi were conducted at 25°C. Scale bars: 10µm. Apical-basal (A-B) is indicated for each close up illustrating an epithelial tissue. In box plots, the black bar indicates the median, the whiskers indicate the maximal range, and each value is indicated by a dot. Statistical tests: T-test in D or Mann-Whitney in C. p-value: *, <0.05; ***, <0.001.

We then tested the impact of increased adherens junction contractility in the hyperplastic tumoral context. To do so, we induced myosin II junctional recruitment through either *flapwing* RNAi or AJ-GFP-trap expression in Yorkie tumours. Strikingly, we observed the apparition of domains containing rounder cells, with delocalised ECad, alpha-Cat, Dlg and Talin. (Fig.3E, F, Sup Fig.4). The cell’s cytoskeleton was also reorganised, with delocalised F-actin, as well as myosin II, around the whole cell cortex, instead of the apical and basal enrichments associated with the epithelial phenotype (Sup Fig.4). Importantly, the conversion of hyperplastic into dysplastic tumours by increased mechanics was fully penetrant, since all manipulated discs were dysplastic, presenting at least one mesenchymal domain (Fig.3G). Hence, while the alteration of myosin II levels at adherens junctions has no impact on the epithelial structure in non-tumoral tissues (Fig.3B, Sup Fig.3B, E), it is sufficient to promote epithelial-to-mesenchymal transition in tumoral discs that normally remain strictly epithelial (Fig.3E, F, Sup Fig.4).

**Figure 4.**
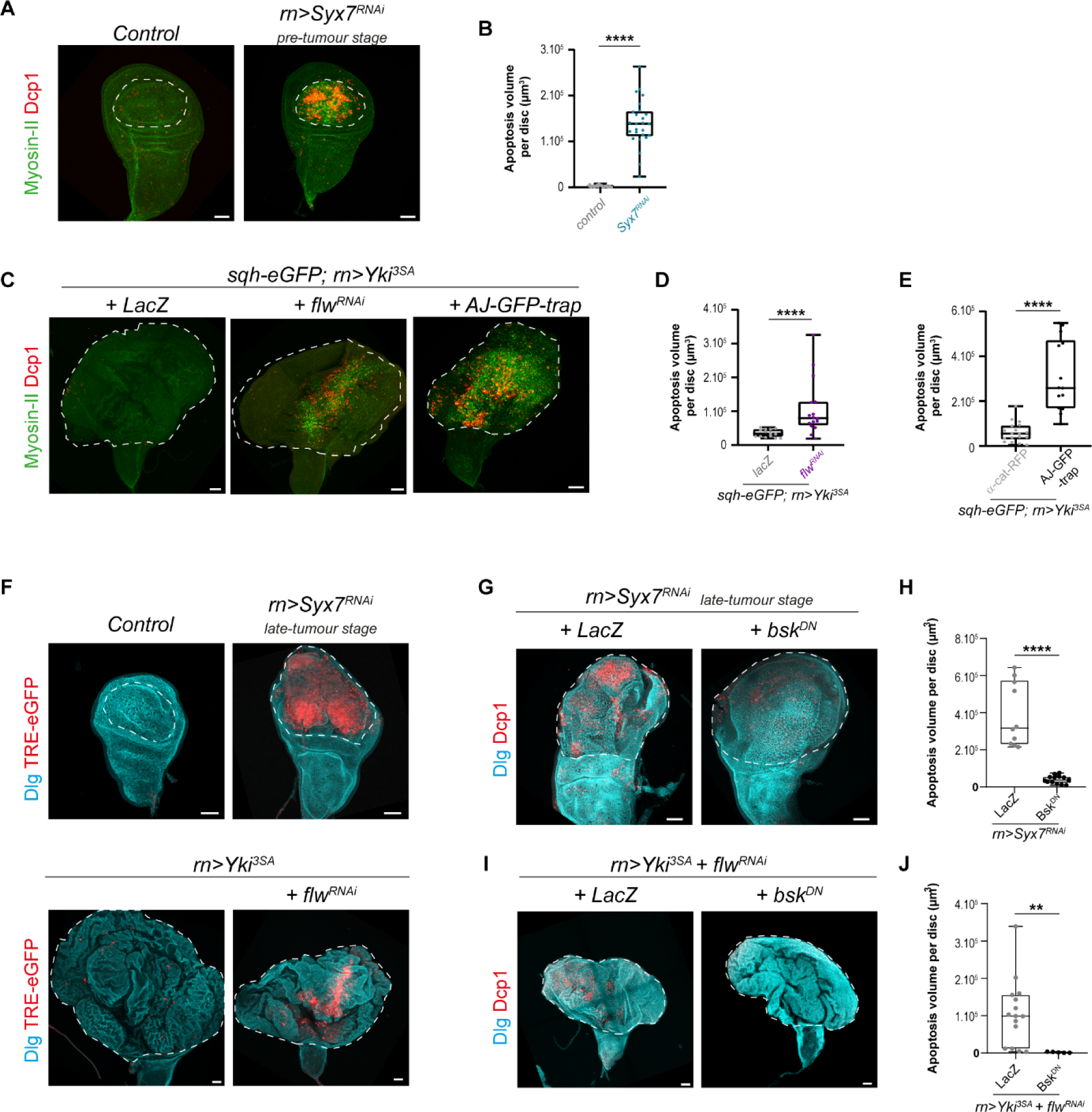
Apoptosis is triggered by high adherens junction tension through JNK activation in tumours. (A) Global views (projections) of pre-tumour stage (96h AEL) wing discs, either control (left) or expressing Syx7 RNAi (right). Note that apoptosis (Dcp-1 staining, red) spatially coincides with the region of myosin II enrichment (green). (B) Quantification of apoptosis in control discs (n=26) and Syx7 RNAi tumours (n=26) at 96h AEL. (C) Global views (projections) of Yorkie tumours (discs overexpressing Yorkie) co-expressing lacZ (used as a control, left), *flapwing* RNAi (middle) or the AJ-GFP trap (GFP trap fused to alpha-Catenin-RFP, right). Note that here again, apoptosis (Dcp-1 staining, red) spatially coincides with regions of myosin II enrichment (green). (D) Quantification of apoptosis in Yorkie tumours concomitantly expressing lacZ (n=14) or *flapwing* RNAi (n=16) to increase myosin II levels. (E) Quantification of apoptosis in Yorkie tumours concomitantly expressing alpha-Catenin-RFP (control, n=18) or the AJ-GFP trap (n=14) to increase myosin II-GFP levels at adherens junctions. (F) Global views (projections) of wing discs of the indicated genotypes (control, n=16; *Syx7* RNAi, n=9; *Yorkie^3SA^*, n=5; *Yorkie^3SA^* + *flapwing* RNAi, n=11). Note that TRE-eGFP expression is detected only in tumours with high myosin II and apoptosis levels (*Syx7* RNAi tumours and *Yorkie* tumours with *flapwing* knockdown). (G-J) Global views (projections) showing the apoptotic pattern (G, I) and associated quantification (H, J) in presence (*lacZ*) or absence (*bsk^DN^*) of JNK signalling driven in *Syx7* RNAi tumours (G, H) or *Yorkie^3SA^* + *flapwing* RNAi tumours. Number of discs analysed: *Syx7* RNAi + *LacZ*, n=10, *Syx* RNAi + *bsk^DN^*, n=13, *Yorkie^3SA^*+ *flapwing* RNAi + *LacZ*, n=15; *Yorkie^3SA^* + *flapwing* RNAi + *bsk^DN^*, n=5. Transgenes are driven in the wing pouch using the *rotund*::Gal4 driver at 25°C, apart from experiments with the AJ-GFP trap at 22°C and with bsk^DN^ (G-J) at 29°C. Scale bars: 50µm. In box plots, the black bar indicates the median, the whiskers indicate the maximal range, and each value is indicated by a dot. Statistical tests: T-test in E or Mann-Whitney in B, D, H, J. p-value: **, <0.018; ****, <0.0001.

### High junctional tension triggers cell death in evolving tumours through the JNK pathway

We next investigated mechanisms that could participate in triggering EMT downstream of increased junctional tension. By looking for alterations that would be present in Syx7 RNAi tumours prior to the onset of EMT, but absent in the control tissue, we observed an important amount of cell death, as revealed by staining against the active form of the effector caspase Dcp1. Indeed, massive apoptosis was observed in pre-tumour Syx7 RNAi discs (96h AEL) in which junctional tension is high (Fig.4A, B), prior to EMT appearing at 108h AEL. Interestingly, apoptosis was absent in Yorkie tumours, apart from a few scattered cells, irrespective of the age of the tumour (Fig.4C, left panel), suggesting that apoptosis was specifically associated with aggressive tumours (see Sup Fig.5A for quantifications). Therefore, in fly tumours, apoptosis is tightly correlated with high myosin II levels, both temporally and spatially, and its appearance precedes the induction of epithelial-to-mesenchymal transition in Syx7 RNAi tumours. Furthermore, Yorkie tumours with increased contractility also presented elevated levels of apoptosis (Fig.4C-E), specifically in regions where myosin levels were higher and transformation occurred (Fig.4C middle and right panels, Sup Fig. 5B). These data further suggest that apoptosis is induced by high level of adherens junction contractility present in evolving tumours.

**Figure 5.**
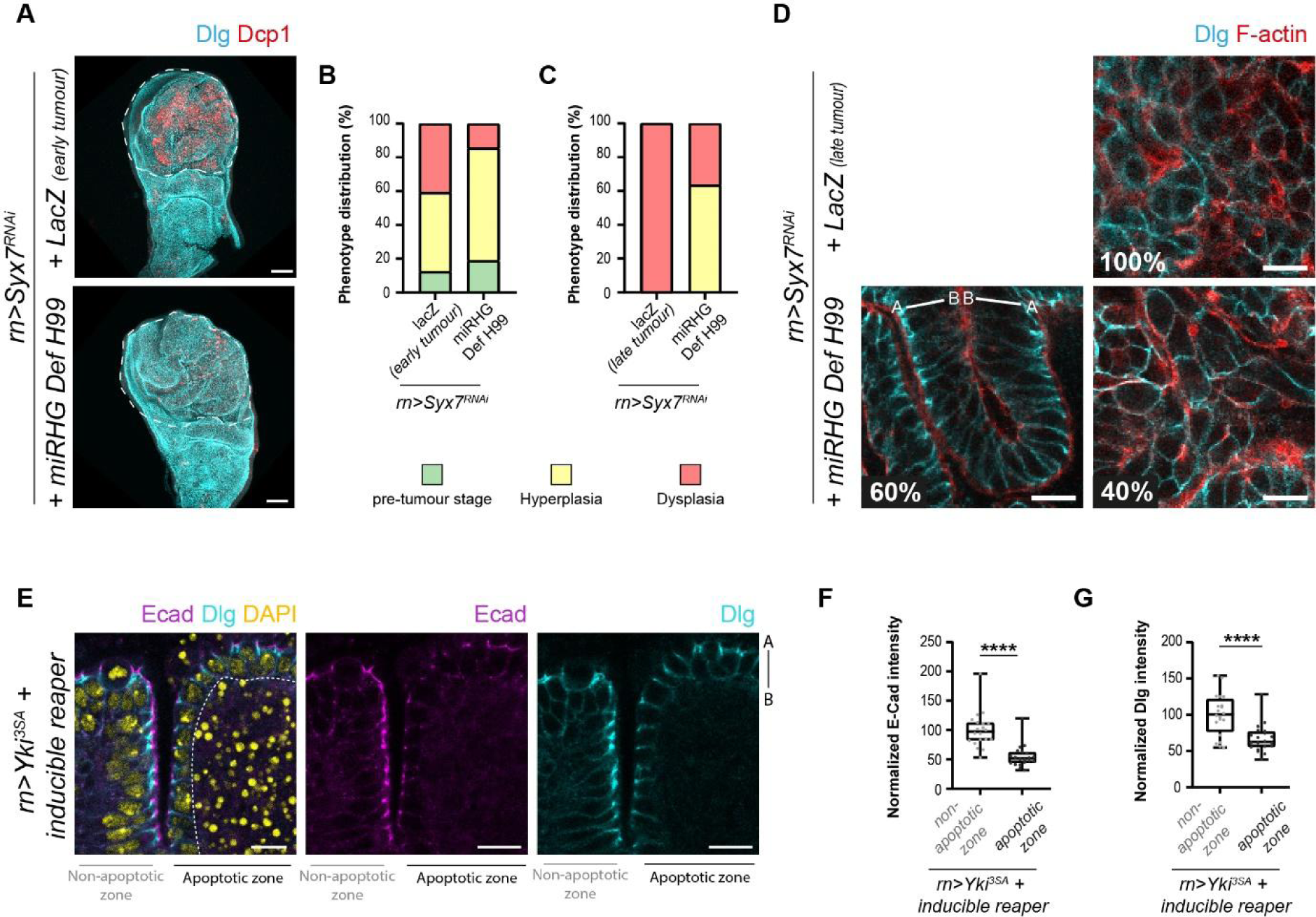
Apoptosis weakens intercellular adhesion and facilitates tumour evolution. (A) Global views (projections) of Syx7 RNAi wing discs of the same age (96h AEL at 30°C, which corresponds to early tumour stage in the control), either co-expressing lacZ (control, top) or miRHG in a Deficiency H99 / + background to inhibit apoptosis (bottom). Note that overall size and shape of Syx7 RNAi tumours with or without apoptosis is similar. (B, C) Histograms of phenotype distribution in Syx7 RNAi discs of the same age with (lacZ) or without (miRHG H99) apoptosis. Control discs are at early tumour stage in (B) (96h at 30°C) and at the late tumour stage in (C) (128h AEL at 30°C). Number of discs analysed: 32 and 19 in lacZ in (B) and (C) respectively; 21 and 22 in miRHG H99 in (B) and (C) respectively. (D) Close up views of Syx7 RNAi tumours of equivalent age with (lacZ) or without (miRHG DefH99) apoptosis, corresponding to tumours analysed in (C). (E) Single plane image showing the state of adherens junctions and septate junctions, respectively labelled with E-Cad (magenta) and Dlg (cyan), in a Yorkie tumour in which apoptosis was induced through controlled *reaper* expression. Apoptosis was massively induced on the right side, but not on the left side of the image. DAPI staining (yellow) labelling nuclei shows the zone of apoptosis induction, with condensed DNA and numerous fragments accumulated. (F, G) Quantification of ECad (F) and Dlg (G) expression levels in apoptotic versus non-apoptotic zones of Yorkie tumours, as shown in (E). Number of discs analysed: 20. Transgenes are driven in the wing pouch using the *rotund*::Gal4 driver at 30°C (panels A-D) or at 25°C (panels E-G). Scale bars: 10µm except 50µm for A. Apical-basal (A-B) is indicated for each close up illustrating an epithelial tissue. In box plots, the black bar indicates the median, the whiskers indicate the maximal range, and each value is indicated by a dot. Statistical tests: Wilcoxon in F and paired T-test in G. p-value: ****, <0.0001.

One candidate to link junctional tension to cell death is the JNK pathway. Indeed, JNK signalling can be activated by Rho1 ^31,32^, a Rho-GTPase which is an upstream regulator of myosin II ^33^, and likely by Myosin II overactivation in *flapwing* mutants ^34^. Moreover, JNK pathway activation is a well-known inducer of apoptosis, especially through the control of the expression of the pro-apoptotic genes such as *rpr* and *hid* ^35–37^.

To investigate the possible regulation of cell death by the JNK pathway, we first analysed the expression of a fluorescent reporter, TRE-eGFP. Its activation was broad in Syx7 RNAi tumours (Fig.4F, top), consistent with previous reports using other reporters of JNK activation such as expression of *puc-lacZ* or MMP1 ^23^. In epithelial Yorkie tumours, TRE-EGFP activation is almost absent, and restricted to a few scattered cells. Interestingly, upregulating myosin II using *flapwing* RNAi in Yorkie tumours led to a massive activation of TRE-eGFP in the center of the disc (Fig.4F, bottom), where both high myosin II and apoptosis are observed.

We next inhibited the pathway at the level of the *Drosophila* JNK, *basket*, using the dominant negative form *bsk^DN^* to determine whether mechanical activation of the JNK pathway triggers tumoral apoptosis. Interestingly, expression of this construct in both Syx7 RNAi tumours (Fig.4G, H) or Yorkie tumours with *flapwing* knockdown (Fig.4I, J) led to a nearly complete loss of apoptosis detection. Together, those results suggest that cell contractility could control apoptosis, a biomarker of early evolving tumours, through the activation of the JNK pathway.

### Apoptosis favours adhesion weakening and tumour aggressiveness

Apoptotic cells are known to influence their surrounding by various signalling or mechanical means during development ^38–40^. Moreover, while apoptosis is a well-known protective mechanism ^1,41^, several reports pointed towards the presence of apoptosis in certain tumoral contexts and its ability to promote cell proliferation in specific cases ^38,42,43^. Together with the close proximity of apoptotic cells to dysplastic zones (Sup Fig. 5B), this prompted us to ask whether apoptosis could contribute to the induction of epithelial-to-mesenchymal transition in *Drosophila* tumours. To investigate the potential pro-tumoral role of cell death, we first inhibited cell death in Syx7 RNAi tumours. Nearly complete cell death inhibition was difficult to obtain, and was only possible through inhibition of the three main fly proapoptotic genes *reaper*, *hid* and *grim* by expression of a RNAi transgene in a genetic background heterozygous for those genes, a genetic combination hereafter referred to as “miRHG DfH99” (Sup Fig.5C-E). We next analysed Syx7 RNAi tumour evolution with or without cell death in tumours grown for exactly the same duration, and which have a similar macroscopic appearance (Fig.5A). Interestingly, when roughly half of the Syx7 RNAi tumours (containing apoptosis) are epithelial and 40% of them are starting to become mesenchymal, Syx7 RNAi tumours without apoptosis presented only 15% of mesenchymal tumours (showing only limited zones of mesenchymal cells), but about 65% of purely epithelial, hyperplastic tumours (Fig.5B). Strikingly, this tendency is amplified at later stages when all Syx7 RNAi tumours bearing apoptosis have turned massively mesenchymal, while blocking cell death maintained 60% of the tumours in an epithelial stage, showing a strong decrease in the number of aggressive tumours at an equivalent time point (Fig.5C, D). These results show that apoptosis promotes the epithelial-to-mesenchymal transition in tumoral contexts.

We next asked how cell death could act as a pro-tumoral factor regarding the epithelial-mesenchymal cell status. To address this question, we first investigated whether hemocytes, the fly macrophages, could play a role. Indeed, macrophages, which are attracted by dead cells to clean apoptotic corpses, are known regulators of tumour aggressiveness in mammals, while hemocytes are indirect modulators of tissue overgrowth when apoptosis is impaired in fly ^44–46^. We observed that some hemocytes are recruited to Syx7 RNAi tumours, but only once they have become dysplastic. Importantly, we did not observe hemocytes in pre-tumoral tissues or in hyperplastic tumours, indicating that their late recruitment could not account for epithelial-to-mesenchymal induction in Syx7 RNAi tumours (Sup Fig.6A, B). We next investigated whether cell death could promote tumour evolution by directly modulating the tension of adherens junctions in Syx7 RNAi tumours, as previously described during development ^39^. Of note, the level of contractility is unchanged by the inhibition of apoptosis, as shown by the fact that apical myosin II levels are similar in the presence or in the absence of apoptosis in pre-tumour Syx7 RNAi discs (Sup Fig.6C-D’). Finally, we investigated whether cell death could directly control the levels of key junctional components whose regulation is important for epithelial-to-mesenchymal transition ^47^. We induced ectopic cell death in the Yorkie tumours that are normally devoid of it and stained tumours with ECad and Dlg to label both adherens and tight junctions. We did not notice epithelial-to-mesenchymal transitions upon massive cell death induction, suggesting that apoptosis *per se* may not be sufficient to induce a complete EMT in tumours. However, we observed that ECad and Dlg junctional levels dropped in epithelial zones with high cell death, compared to epithelial zones with low or no cell death (Fig.5E-G). Quantifications show that Dlg and ECad levels in living cells are decreased by 33% or 45% respectively in presence of high apoptosis levels nearby, when compared to epithelial zones with low or no cell death in close vicinity (Fig.5F, G). This indicates that apoptosis non-autonomously weakens cell-cell junctions in tumours. Altogether, this led us to propose that apoptosis acts downstream of increased contractility of tumour cells to facilitate EMT and therefore tumour evolution, through the destabilization of both adherens and tight junctions.

**Figure 6:**
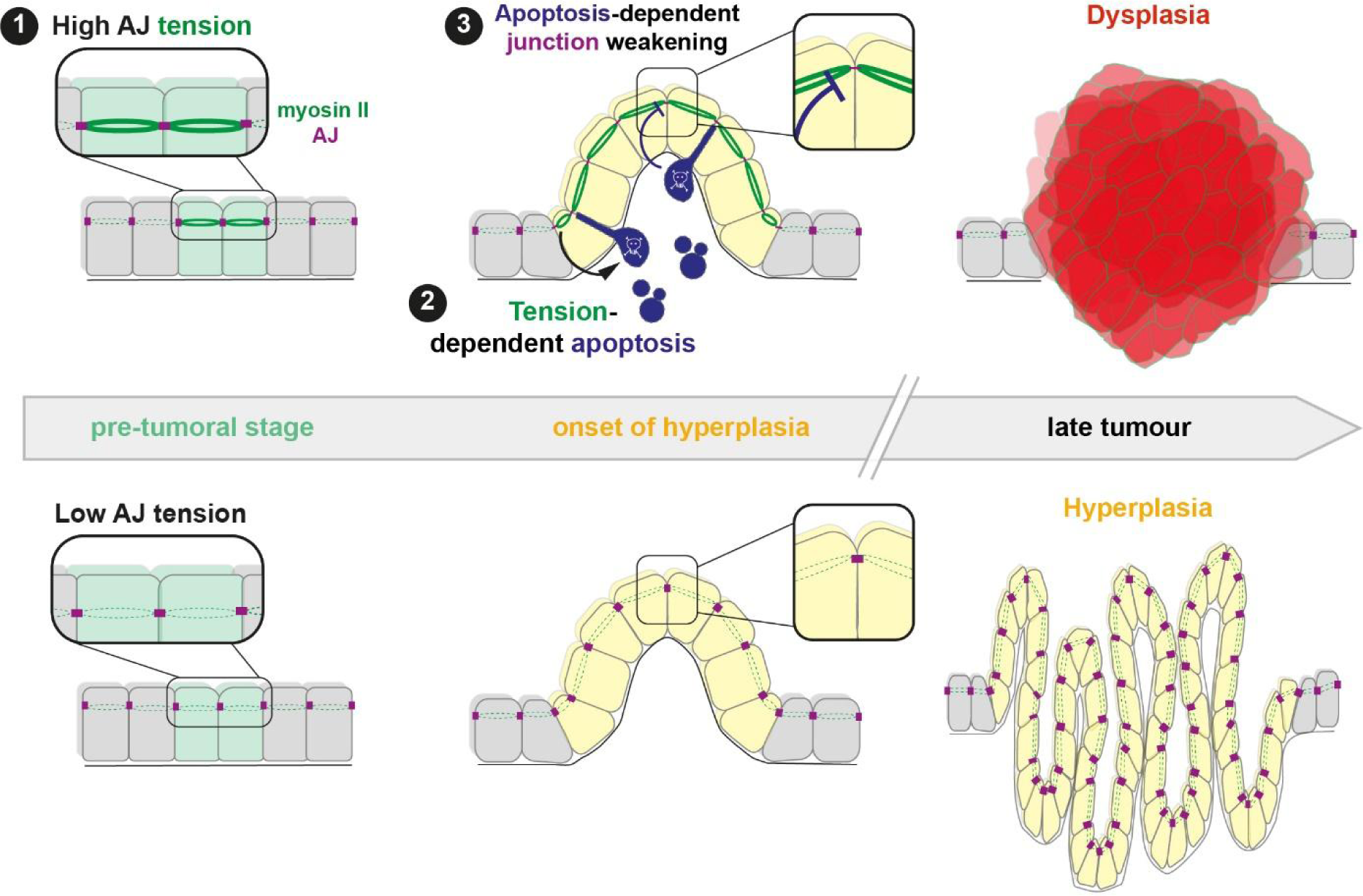
Model of tumour evolution controlled by adherens junction tension and apoptosis. Tumour evolution from the pre-tumoral stage (left) to late tumours (right) depending on whether adherens junctions (AJ, magenta) are under high tension (top row) or low tension (bottom row). High adherens junction tension (1) is caused by myosin II accumulation (dark green) starts at the pre-tumoral stage (cells in light green) and induces apoptosis (2). Apoptosis (blue) triggers weakening of adherens junctions (3) and tight junctions (not illustrated) in surrounding living cells, thereby facilitating the switch from hyperplasia (yellow cells) to dysplasia (red cells). High AJ tension can also triggers dysplasia, independently of apoptosis (not depicted). On the contrary, when tension is low at AJ, hyperplasia expands but does not switch into dysplasia.

## DISCUSSION

So far, alterations of tumour mechanical properties, such as increased extracellular matrix rigidity, increased pressure or compressive growth, have been essentially documented at late tumour stages, or in cultured cells which are not in a physiological environment. Here our results show that early during tumour development, high junction contractility induces the presence of massive apoptosis in a JNK-dependent manner to eventually favor the shift from hyperplastic to dysplastic tumour stages through the non-autonomous weakening of adherens and tight junctions (Fig.6, top panel), while low-contractility tumours remain hyperplastic (Fig.6, lower panel). Overall, our results identify the intrinsic mechanical properties of pre-tumoral tissues as a trigger of tumour aggressiveness in an *in vivo* model system.

A key feature of tumour evolution is the activation of the EMT program. EMT is well known to be an inducible change in cell behaviour, controlled by intercellular signalling pathways such as TGFbeta ^47^. The tumour microenvironment, through signalling from cancer associated fibroblasts (CAFs) or tumour associated macrophages (TAMs) or remodelling of the surrounding ECM is associated with tumoral EMT regulation ^8,48^. Yet, our results point towards an unsuspected alternative pathway to trigger EMT, which is autonomously controlled by tumour cell mechanics. Although adherens junctions are known to be mechanosensitive in epithelial tissues, this work identifies them as key mechanosensitive structures driving tumour evolution. During development, endocytosis can be enhanced at adherens junctions under increased tension ^49^, and mechanosensitive endocytosis was shown to control E-Cad and membrane removal to maintain junction homeostasis ^50^. Forces can also impact tight junctions through ZO-1 regulation in cultured cells ^51^. However, as we report for fly tissues, enhanced junctional tension is not sufficient to trigger EMT during development, as it does in tumours. This suggests a shift in the mechanosensitive properties of cells in pre-tumoral tissues, which will necessitate further investigations in the future.

A striking result is that high adherens junction tension induces cell death which favours dysplasia formation. This points towards a far more complex role of apoptosis than previously anticipated in tumours biology. While increased resistance to apoptosis is long considered as a hallmark of cancer ^52^, more recent publications hinted at a paradoxical pro-tumoral role by promoting cell proliferation ^42,43,53^, while failed apoptosis would favour invasiveness ^54^. Our data further show that, by fragilizing intercellular adhesions, cell death could also potentiate EMT. Given that mesenchymal tumour cells are associated with increased resistance to treatments and acquisition of cancer stem cell (CSC) traits ^9,11^, massive but incomplete cell death induction by radiotherapy or chemotherapy, depending on specific tumour properties (such as the state of tumour cells mechanics) could favour the appearance of pockets of more resistant cells and, on the long run, tumour repopulation.

Finally, the impact of mechanics on tumour biology takes place at very early stages of tumour formation, when overgrowth has not yet taken place. With those results in mind, it would be interesting to revisit tumour systems in other model organisms in which circulating EMT cells are detected before tumours could be histologically identified, as is the case in the pancreas ^55^. In that case, alterations in the junctional myosin II pattern and associated tension could not only induce EMT but also be an early biomarker of aggressiveness.

## Acknowledgments

We thank Alice Davy and Corinne Benassayag for their constructive comments on the manuscript. We thank K.Sugimura, J.H.Park, H.Steller, P.Léopold and BDSC for sharing fly stocks and J.Zallen and DGRC for providing reagents.

## Author contributions

MM performed the experiments with the help of BM. BM and MS supervised the project and wrote the manuscript. MS provided funding.

## Competing interests

Authors declare that they have no competing interests.

## Funding

MS’s lab is supported by grants from the National Agency of Research (ANR, PRC AAPG2021, CellPhy) and from the Research Association against Cancer (ARC, Programme Labélisé AAP2020, ARCPGA12020010001154_1591). MM was supported by MENRT and ARC fellowships.

## Data and materials availability

All data are available in the main text or the supplementary materials.

## Methods

### Fly stocks

Knock-in and protein trap fluorescent proteins available from Bloomington Drosophila Stock Center (BDSC) are: ECad-GFP (BDSC_60584); rhea[MI00296]-mCherry/TM6B (to reveal the Talin protein, BDSC_39648), Baz[CC01941]-GFP (BDSC_51572), dlg1[MI06353-GFSTF.0] (BDSC_59417) and TRE-eGFP (BDSC_59010). The sqh-eGFP[29] was used to reveal Myosin II distribution and was described previously ^28^. Viking-TagRFPt is a knock-in designed and generated by homologous recombination by InDroso functional genomics (Rennes, France). The tag was inserted in N-terminal just before the ATG, and the resulting flies were validated by sequencing.

Drivers used in this study available from Bloomington Drosophila Stock Center (BDSC) are: rn::Gal4 (BDSC_7405); pdm2::Gal4[GMR11F02] (BDSC_49828) and Hml::QF2.Delta (BDSC_66468).

Responsive UAS or QUAS lines obtained from BDSC were: UAS::nls-GFP (BDSC_4776); UAS::RNAi-flw[HMS04465] (BDSC_57022); UAS::Yorkie[3SA] (BDSC_28817); UAS::Diap1-H (BDSC_6657), UAS::bsk.DN (BDSC_6409) and QUAS::6x-mCherry-HA (BDSC_52270). A UAS::lacZ[j] stock was generated in this study by backcrossing the UAS::lacZ.Exel line (BDSC_8529) obtained from BDSC with *w*^1118^ flies. The UAS::Vhh4-alpha-cat-TagRFPt stock (i.e. AJ-GFP trap) was described previously in ^26^ and the HS::Flp on the X chromosome was reported in ^39^.

The following stocks were gifts: UAS::alpha-cat-TagRFP stock was obtained from K. Sugimura (iCeMS, Japan), UAS::miRHG from JH. Park (University of Tennessee, USA) and UAS::FRTstopFRTreaper[10.3] from D. Fox (Duke University, USA). The Deficiency Df(3L)H99 (BDSC_1576) that removes the three pro-apoptotic genes *hid*, *reaper* and *grim* was a gift from H. Steller (Rockefeller University, USA).

w; elav::Gal80; rn::Gal4/TM6B and w; elav::Gal80; rn::Gal4, UAS::RNAi-Syx7[GD2767]/TM6B, tub::Gal80 stocks were gifts from P. Léopold (Curie Institute, France). The UAS::RNAi-Syx7[GD2767] original line was originally generated at Vienna Drosophila Resource Center (VDRC). The w; rn::Gal4, UAS::Yorkie[3SA]/TM6B, tub::Gal80 stock was generated in this study. Maintenance of transgenes on the second chromosome in these backgrounds was achieved by using a red fluorescently labelled CyO balancer (BDSC_35523).

Experiments were performed at 25 degrees in both males and females indifferently, unless indicated (see below). Loss of function experiments using Syx7 RNAi were carried out at 25 degrees or 30 degrees, depending respectively on the absence or the presence of an additional UAS transgene. Experiments involving the AJ-GFP trap transgene, either during development or within Yorkie tumours, were conducted in males only, at 21 degree, to avoid too high induction of cell death.

### Immunostainings and image acquisition

Wing discs at the indicated stage (see below) were dissected in PBS 1x. Tissue were fixed by paraformaldehyde 4% diluted in PBS 1x during 20 minutes. Then the samples were washed and saturated in PBS 1x, 0.3% triton x-100 and BSA 1% (BBT). Next, the samples were incubated overnight at 4°C with primary antibodies diluted in BBT. Samples were washed for 1h in BBT before a 2h incubation at room temperature with secondary antibodies diluted in BBT. Finally, samples were washed with PBS 1x, 0.3% Triton x-100 for 1h and mounted in Vectashield containing DAPI or TRITC-Phalloidin (Vector Laboratories). A 120-mm deep spacer (Secure-Seal^TM^ from Sigma-Aldrich) was placed in between the glass slide and the coverslip to preserve morphology of the tissues.

The actin cytoskeleton was labelled using the Rhodamine-Phalloidin probe from Fischer Scientific (1/500, 2h incubation together with secondary antibodies). Alternatively, samples were mounted in Vectashield hardset medium complemented with TRITC-Phalloidin (Vector Laboratories). Nuclei were labelled with DAPI included in Vectashield mounting medium, when necessary.

Primary antibodies obtained from Developmental Studies Hybridoma Bank (DSHB) were: anti-Dlg (4F3-c, mouse, 1/200); anti-ECad (DCAD2-c, rat, 1/100); anti-betaPS (CF.6G11, mouse, 1/50). Anti-cleaved Dcp1 (#9578, rabbit, 1/200) was purchased from Cell Signaling Technologies. Anti-Baz (guinea pig, 1/500) was a gift from J. Zallen (Sloan Kettering Institute, USA).

Secondary antibodies (Alexa-488, −555 and −647) were purchased from Interchim and diluted at 1:200, except Alexa-647 at 1:100.

Fixed samples were imaged using inverted Zeiss laser scanning confocal microscopes, either LSM880 or LSM900, 25x/0.8 multi immersion, 40x/1.4 oil or 63x/1.4 oil objectives and equipped with a piezo stage. Images were processed with the Fiji and Imaris softwares.

### Kinetics of tumour development

To perform kinetics of tumour development, or compare tumours at a given timepoint, the following strategy was used. Virgin females (ultimately 250-300 per cage) were collected in 3-5 days. Males were crossed with females (about 10 males for 25-30 females) in tubes. Crosses were transferred into cages to lay eggs on standard fly food medium. Egg-laying was performed for 3 hours at 25°C. Petri dishes containing the fly food medium and the eggs was transferred either at 25°C (when UAS::Syx7 RNAi is the sole UAS transgene) or at 29°C (when we used second UAS transgene in addition to the UAS::Syx7 RNAi). On the next day, the medium with eggs was transferred into standard plastic vials also containing standard fly food, and replaced at 25°C or 29°C. Eggs were aged for the indicated time before dissection, younger discs being obtained after 96h of incubation at 25°C (i.e. 96h after egg laying, AEL) or 64h of incubation at 29°C. Timepoints analysed were separated by 12 or 24h, with a temporal resolution of 3hours (corresponding to the length of the egg-laying). For each experiment, control crosses were performed in parallel by replacing the UAS::miRHG, UAS::Diap1 or UAS::flapwing RNAi by a neutral UAS::lacZ transgene or the UAS::Vhh4-alpha-cat-TagRFPt (AJ-GFP-Trap) by the UAS::alpha-cat-TagRFP.

Note that control discs could not be analysed past 120h AEL. This is because control individuals enter metamorphosis at this timepoint, and wing discs start to undergo morphogenesis, precluding further comparison with tumoral discs. To determine their size, tumoral discs of 144h AEL and beyond were therefore compared to 120h AEL control discs.

### Classification of tumour phenotypes

In our experiments, wing discs were classified in 3 categories: normal, hyperplastic and dysplastic. This subdivision takes 2 parameters into account, the overall size of the wing pouch and the epithelial vs mesenchymal organization of the tissue within the pouch.

For such an analysis, we systematically generated control (non-tumoral discs in which UAS::Syx7 RNAi or UAS::Yorkie[3SA] were replaced by a neutral UAS::lacZ transgene) of the same age. This allowed to determine whether the pouch was overgrown or not.

We also stained the discs with a combination of markers of epithelium organization: ECadherin or alpha-Catenin to label adherens junctions, Disc Large (Dlg) to label septate junctions, beta-Integrin or Talin to label basal adhesions, or Bazooka/Par3 as a marker or apico-basal polarity. Both fluorescent knock-in/protein trap lines and antibodies were used. A combination of DAPI/Phalloidin was also used, alone or in combination with the above markers. Organization of the tissue in a monolayer structure, with defined apical and basal poles, was classified as epithelial. In mesenchymal zones, the monolayer structure is lost, producing mass-like structures in which apico-basal polarity is lost. This is reflected for example by the proximity of apical and basal markers, the F-actin cytoskeleton that surrounds the whole cell or the spacing between nuclei which increases in mesenchymal zones.

Experimental wing pouches which were of a similar size than control pouches at the same timepoint, and in which the tissue is strictly epithelial were classified as “normal”. Experimental wing pouches wider than the pouch of control discs, and in which the tissue is strictly epithelial were classified as “hyperplastic”. Finally, experimental wing pouches wider than the pouch of control discs, and in which at least one zone is organized in a mesenchymal-like fashion were classified as “dysplastic”. Data were represented as 100% stacked histograms, with normal, hyperplastic and dysplastic discs coloured in green, yellow and red respectively. Number of discs analysed in indicated in figure legends.

### Apoptosis ectopic induction

Induction of apoptosis was performed by driving an inducible UAS::FRTstopFRT-reaper transgene together with UAS::Yorkie[3SA] in the whole wing pouch. Crosses were performed and larvae initially developed at 25°C, producing hyperplastic tumours. A heat shock was performed at 37°C for 20 minutes, and larvae were grown for 16 additional hours at 25°C before dissection.

### Laser dissection

Tissue culture was performed as follow: wing discs were dissected in Schneider’s insect medium (Sigma-Aldrich) supplemented with 15% fetal calf serum and 0.5% penicillin-streptomycin. Wing discs were transferred on a glass slide in 13.5 uL of this medium confined in a 120 mm-deep double-sided adhesive spacer (Secure-Seal^TM^ from Sigma-Aldrich). A high precision glass coverslip (n°1.5H, Marienfeld) was then placed on top of the spacer. Dissection tools are cleaned with ethanol before dissection.

Laser dissection experiments were performed on a Zeiss LSM880 laser scanning microscope fitted with a pulsed DPSS laser (532 nm, pulse length 1.5 ns, repetition rate up to 1 kHz, 3.5 µJ/pulse) steered by a galvanometer-based laser scanning device (DPSS-532 and UGA-42, from Rapp OptoElectronic, Hamburg, Germany). The laser beam was focused through a water-immersion lens of high numerical aperture (Plan-Apochromat 63x from Zeiss). Experiments were performed using a numerical 2x zoom. Photo-disruption at adherens junctions was produced in the focal plane by illuminating at 80 % laser power a 20 pixel line (3 runs). Images of ECadherin-GFP were acquired every 500 ms using a 488 nm Argon laser and a GaAsP photomultiplier.

### Statistical analysis

Statistics were performed in Prism. N and p values are indicated in figure legends. Box plot were generated in Prism and represent the median, 10 and 90 percentiles.

The normality of data sets was determined using Prism 8 (GraphPad). For data sets that follow a normal law, a two-tailed unpaired Student’s t-test was used to assess the significance. Otherwise, a Mann Whitney test was performed.

Significance is denoted as follows according to the p value: ****p<0.0001; ***p<0.001; **p<0.01; *p<0.05; NS p>=0.05 (not significant). Box plot were generated with Prism.

### Quantifications

#### -Acto-myosin and Collagen IV intensity

To quantify acto-myosin, confocal Z-stacks were acquired, with a 25x multi immersion objective, and a resolution of 1.11µm in x/y and 1µm in z respectively. We systematically imaged the whole disc, both in x-y and in depth. Z-stacks were analyzed by generating sagittal sections with the Fiji software. For each sagittal section, two segmented lines (with a thickness of 5) were drawn on the apical surface of the epithelium: one in the notum/hinge region (internal control) and one in the pouch region (experimental zone). The mean fluorescence intensity for each marker of interest was measured and the value in the pouch was divided by the value in the unmanipulated notum/hinge region to obtain a normalized value. 3 sagittal sections were analyzed with this strategy per disc, and the averaged value was calculated. Analysis was conducted on tissues stained with Phalloidin to reveal the F-actin cytoskeleton, and on the sqh-eGFP knock-in line to visualize myosin II.

For quantifications on the basal side, a similar strategy was performed, except segmented lines with a thickness of 3 were used. Quantifications of Collagen IV were performed on the Viking-TagRFPt knock-in line, with the segmented lines drawn on the basal side of the epithelium by following Viking-TagRFPt staining. For quantification of basal stainings, the control region analyzed was the hinge region, in with there is only a monolayer epithelium, as in the pouch. The notum region was avoided, since a layer of mesenchymal-like adepithelial cells lies underneath the disc epithelium, which could complicate Collagen IV measurements.

#### -ECadherin and Disc Large intensity after cell death induction

Quantifications of E-cadherin and Disc large after cell death induction were performed on tissues obtained as indicated in the “Apoptosis ectopic induction” section. Single plane confocal images were acquired with 40x/1.4 oil objective, with a x/y resolution of 0.078 µm and an image size of 2048*2048 pixels. Images were taken in region in which a zone with massive apoptosis was in close vicinity to a zone with low/no apoptosis. Segmented lines were drawn at the level of adherens junctions (ECad staining, line thickness of 15) or at the level of septate junctions (Dlg staining, line thickness of 25). 2 to 3 measures were made in both the apoptotic and the non-apoptotic zones, which were recognized based on nuclei integrity.

#### -apoptosis volume

To quantify apoptosis levels, discs were stained with the anti-cleaved Dcp1 antibody, which labels the activated form of the executioner Caspase Dcp1. Discs were counterstained with DAPI to visualize the whole disc morphology. Confocal Z-stacks were acquired, with a 25x multi immersion objective. We systematically imaged the whole disc, both in x-y and in depth. 3D image stacks were reconstructed using Imaris software (Bitplane). The channel corresponding to the apoptotic staining was segmented in Imaris and the total volume was calculated by summing up the volume of each apoptotic cell or fragment.

#### -macrophages number

A similar strategy than the one used to quantify apoptosis volume was used, except that the Spot detection function was used in Imaris. A threshold of 7µm was set up to detect the spot, based on the average size of microphages initially determined. Macrophages were labelled using expression of a QUAS::mCherry fluorescent protein driven by the Hml::QF2 driver.

## Notes

### Competing Interest Statement

The authors have declared no competing interest.

