## Supplemental figures for "Mechanical state of pre-tumoral epithelia dictates subsequent tumour aggressiveness"

Supplementary Materials

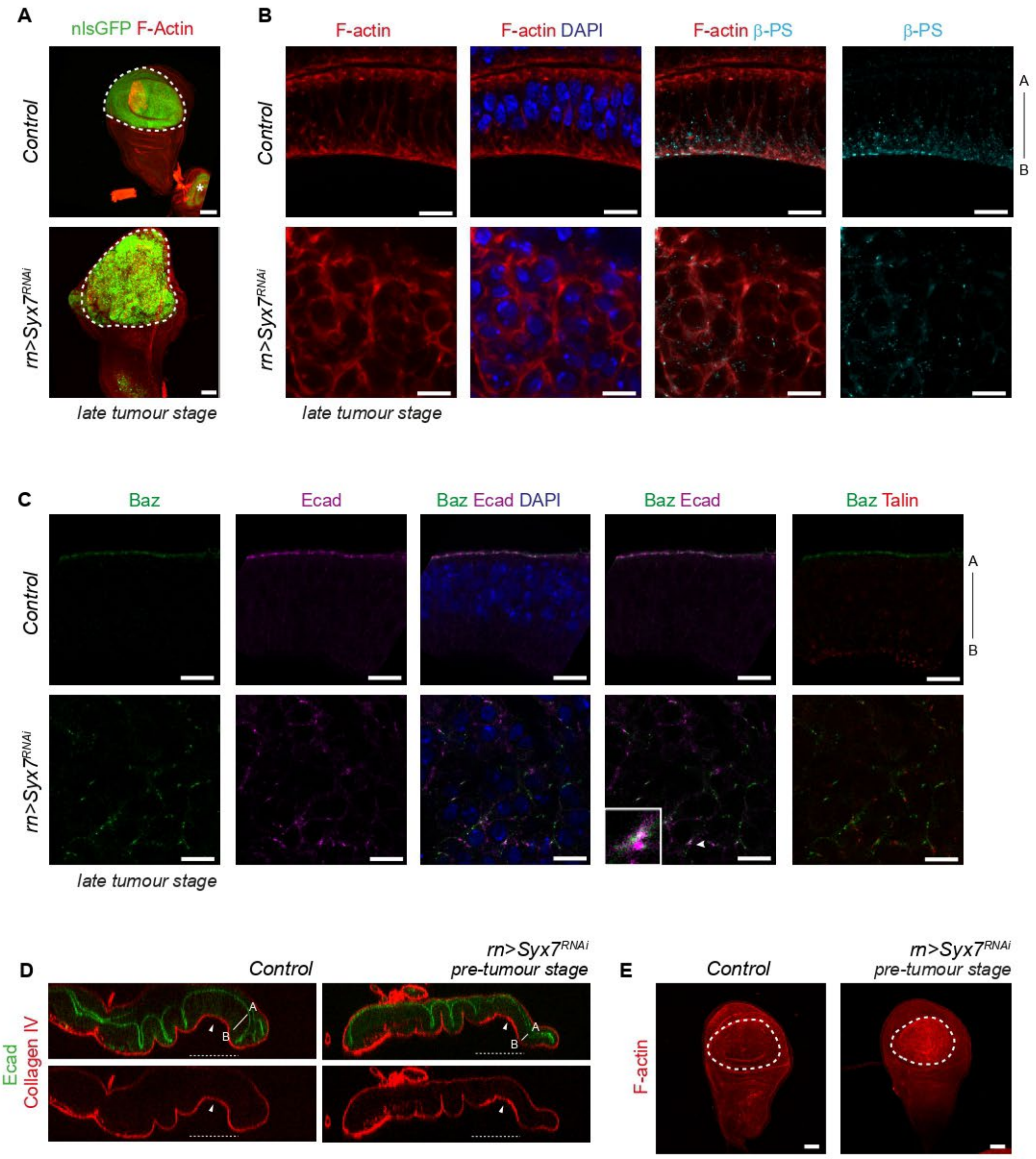


**Supplementary Fig. 1**

**Supplementary Figure 1. Additional characterization of Syx7 RNAi tumour organisation**

(A) General views (projections) of *Drosophila* wing discs stained for F-actin (red) to reveal overall tissue morphology in the control (top) or Syx7 RNAi disc at the late tumour stage (bottom). A nuclear GFP (green) is expressed in the wing pouch to highlight the domain of interest in this study (surrounded by a dashed line). Note that the green staining indicated by the asterisk in the control correspond to a haltere disc, which is not analysed in this study. (B, C) Close up views showing tissue organisation in control discs (approximately 120h AEL, top row) and late tumour stage Syx7 RNAi discs (approximately 144h AEL, bottom row). In (B), discs were stained with phalloidin to reveal F-actin and beta-PS antibody to reveal Integrin receptors. Note tissue disorganisation associated with epithelial-to-mesenchymal transition in Syx7 RNAi tumours, with rounder cells instead of a columnar organisation, F-actin redistributing al around the cell cortex instead of accumulating essentially at apical and basal poles, and mislocalised dots of Integrins. In (C), discs were labelled using endogenous E-Cadherin fused to GFP (ECad, magenta), endogenous Talin fused to mCherry to label Integrin associated basal adhesions (red), and an antibody against the junctional component Par3/Bazooka (Baz, green) which is a key component of epithelial apico-basal cell polarity. Like in (B), note the disorganisation of the tissue following Syx7 knockdown, with mislocalised dots for each marker. Dots for apical (ECad, Baz) and basal (Talin) markers can be observed in close vicinity, indicating the loss of polarity. Note also that while Ecad and Baz colocalize in control epithelium, they do not systematically colocalize in Syx7 RNAi tumours (see inset taken at the level of the arrowhead). Number of discs analysed are respectively 6 and 2 in control and Syx7 RNAi in (B) and 11 and 9 in control and Syx7 RNAi in (C). (D) Analysis of extracellular matrix distribution in control (left, n=15) and Syx7 RNAi (right, n=12) at the pre-tumour stage (96h AEL) in the wing pouch (underlined by dashed lines). Representative sagittal sections are shown, with high ECM staining on the basal side (arrowhead), as revealed by Collagen IV-TagRFPt staining (red). The epithelium, and especially its apical side, is revealed by ECad staining (green). At that stage, Collagen IV is normally distributed in the wing pouch of Syx7 RNAi tumours. (E) General views (projections) of *Drosophila* wing discs stained for F-actin (red) to reveal overall tissue morphology in the control (left) or Syx7 RNAi disc at the early tumour stage (right). The wing pouch is indicated by a white dashed line. Scale bars: 50µm in A and E, 10µm in B and C. A-B indicate apical-basal polarity in the control and Syx7 RNAi pre-tumoral epithelium.

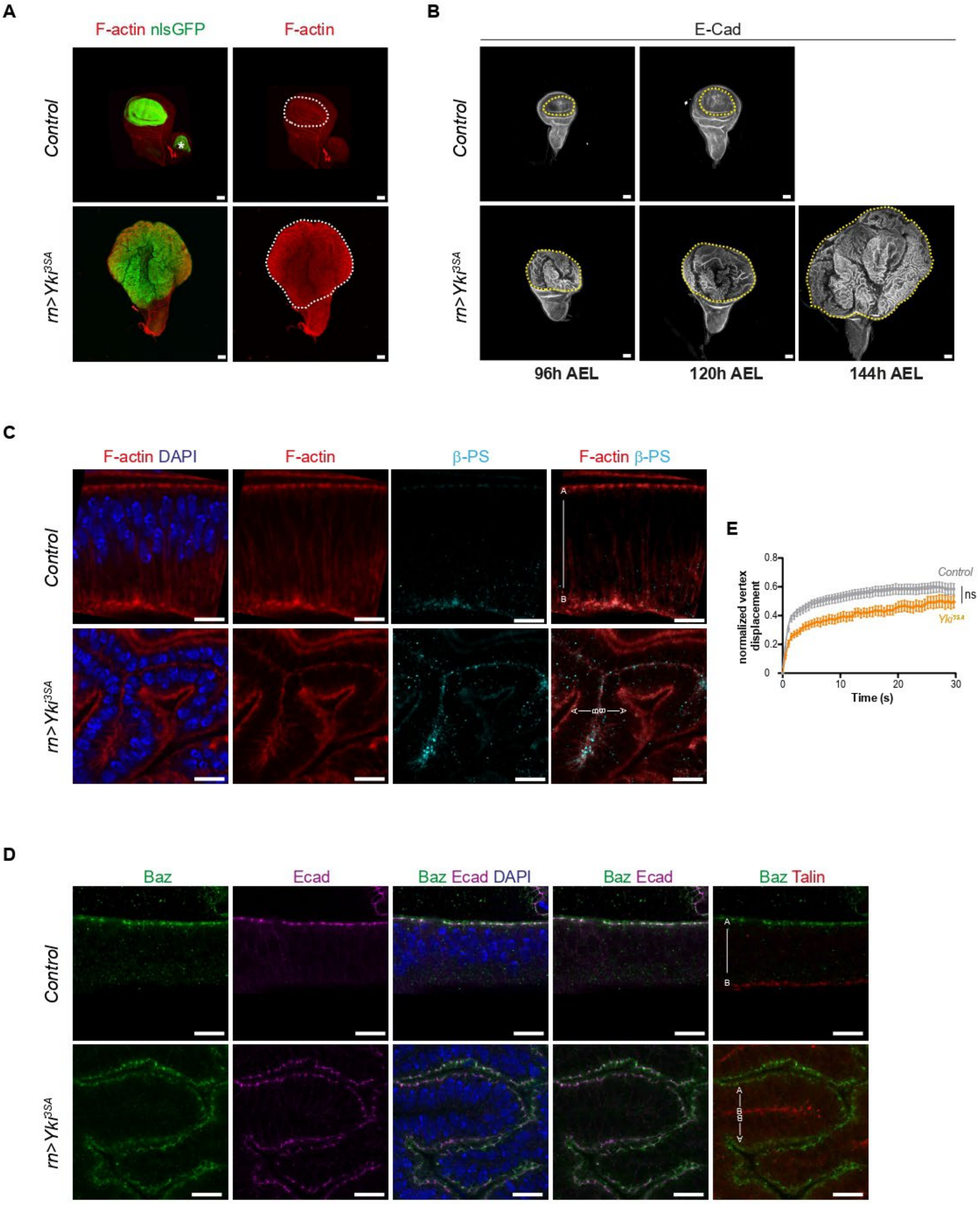


**Supplementary Fig. 2**

**Supplementary Figure 2. Additional characterization of Yorkie tumours**

(A) General views (projections) of *Drosophila* wing discs stained for F-actin (red), to reveal overall disc morphology, and expressing a nuclear GFP in the wing pouch to reveal its size in control (top) and following Yorkie overexpression (bottom). (B) General views (projections) of *Drosophila* wing discs stained for E-Cadherin (white) to reveal overall tissue morphology at the indicated timepoints (+/- 1.5 hour) for control (top row) or Yorkie overexpression (bottom row) conditions. The wing pouch is delineated by the yellow dashed line. Note the strong increase in pouch size in discs overexpressing Yorkie compared to the control at any timepoint as well as at successive timepoints. Control animals enter pupariation just after 120h AEL, preventing further analysis of larval tissues in this condition. On the contrary, Yorkie tumours lead to an extension of larval life, and further analysis of larval tissues is possible at timepoints such as 144h AEL and beyond. Number of discs analysed are indicated in legend of Figure.2J. (C, D) Close up views showing tissue organisation in control discs (approximately 120h AEL, top row) and late Yorkie tumours (approximately 144h AEL, bottom row). DAPI was used to reveal nuclei (blue). A-B indicate apical-basal polarity. In (C), discs were stained with phalloidin to reveal F-actin, and beta-PS antibody to reveal Integrin receptors. Note that despite multiple convolutions, the tissue keeps an epithelial organisation in Yorkie tumours, with a columnar organisation, F-actin still accumulated essentially at apical and basal poles, and strictly basal dots of Integrins are observed. In (D), discs were labelled using endogenous E-Cadherin (ECad) fused to GFP (magenta), endogenous Talin fused to mCherry to label Integrin associated basal adhesions (red), and an antibody against the junctional component Par3/Bazooka (Baz, green) which is a key component of epithelial apico-basal cell polarity. Like in (C), note the polarized organisation of the tissue following Yorkie overexpression, and the colocalization of ECad and Baz, as expected for epithelial cells. Apical (ECad, Baz) and basal (Talin) markers are observed at opposite cell poles, indicating the maintenance of epithelial polarity. Number of discs analysed are respectively 6 and 8 in control and Yorkie overexpression conditions in (C) and 11 and 14 in control and Yorkie overexpression conditions in (D). (E) Analysis of adherens junction tension by laser ablation in control (grey) and Yorkie (orange) discs expressing E-Cadherin-GFP at 96h AEL. Curves of vertex displacement following laser ablation are shown. The number of junctions analysed in control and Yorkie overexpression conditions are respectively of 33 and 12. Related to Fig.2L. Scale bars: 50µm. in A, B and 10µm in C, D. Statistical test: T-test in E. p-value: ns, non-significant.

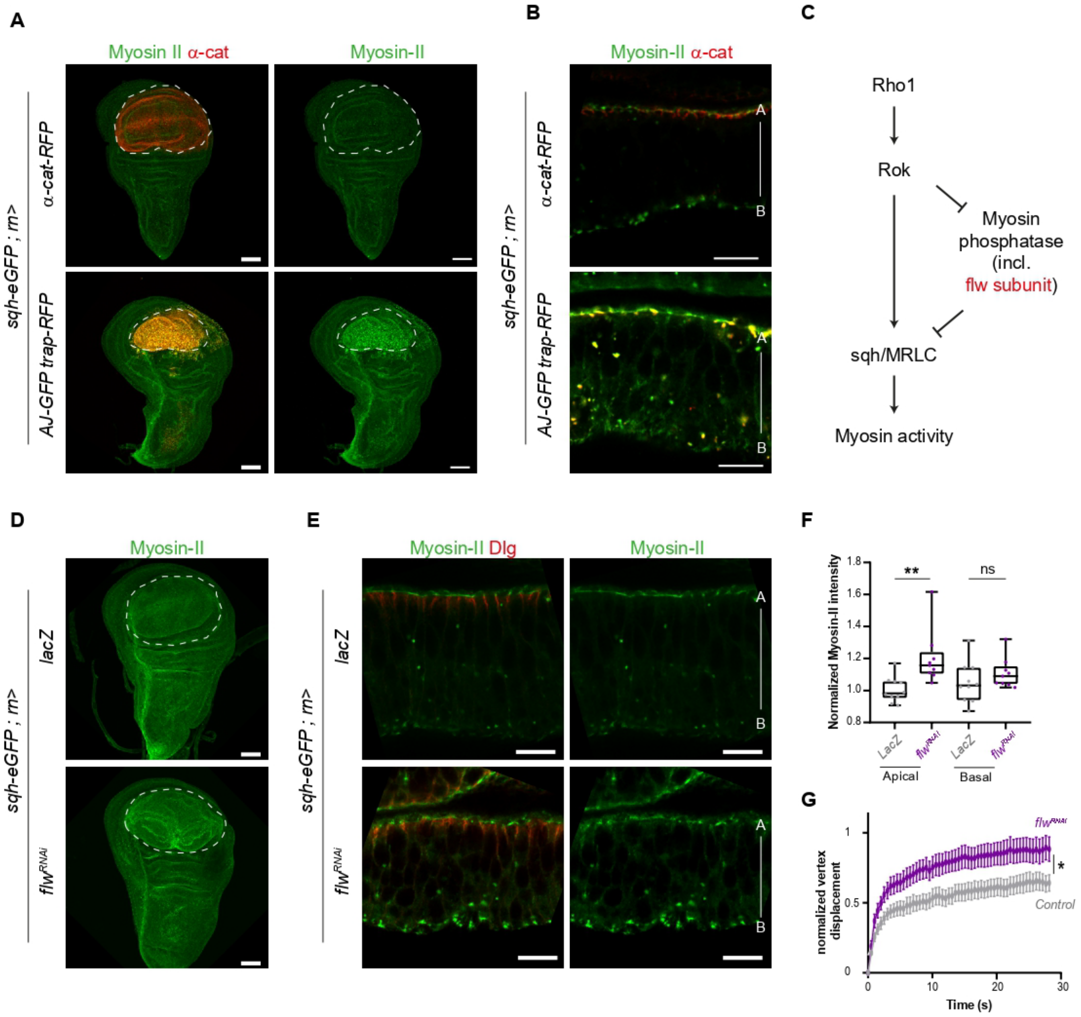


**Supplementary Fig. 3**

**Supplementary Figure 3. Increase in myosin II levels at adherens junctions caused by flapwing inhibition or AJ-GFP-trap expression**

(A, B) General views (projections) and close up views (single plane, B) of wing discs in which either alpha-catenin-TagRFP (control, top) or the AJ-GFP-trap (bottom) are expressed specifically in the pouch (indicated by the dashed line). The AJ-GFP-trap is constituted by a fusion between a nanobody that recognizes GFP, the adherens junction constituent alpha-Catenin and the TagRFP fluorescent molecule to follow its subcellular localisation. The regulatory subunit of myosin II, Sqh, is endogenously tagged with eGFP. (B) The close-up views illustrate that myosin II (green) levels are increased at the level of adherens junctions in presence of the AJ-GFP-trap (colocalization with the TagRFP, red). Note that myosin II alone and Dlg staining from the same images are shown in Fig.3B. (C) Scheme of myosin II regulation by a balance of phosphorylation/dephosphorylation. (D, E) General views (projections, D) and close up views (single plane, E) of wing discs either control (expressing lacZ, top) or in which the myosin II phosphatase activity is reduced due to flapwing RNAi expression (bottom). The wing pouch is indicated by a dashed line. Note the increase in myosin II levels caused by flapwing inactivation. (E) The close-up views illustrate that the apico-basal polarity is maintained, with Dlg (red) still localized at apical tight (septate) junctions. Number of discs analysed: 5 and 7 in control and *flapwing* RNAi respectively. (F) Quantification of myosin II-GFP levels at adherens junctions (apical) or basal junctions (basal) in control (expressing lacZ, n=10) and flapwing RNAi wing discs (n=9). (G) Analysis of adherens junction tension by laser ablation in control (expressing lacZ, grey) and flapwing RNAi discs (magenta) co-expressing E-Cadherin-GFP. Curves of vertex displacement following laser ablation. Number of junctions analysed: 14 and 16 in control and flapwing RNAi respectively. Related to Fig.3D.

Transgenes are driven in the wing pouch using the *rotund*::Gal4 driver at 25°C, apart from experiments with the AJ-GFP trap which were conducted at 22°C. Scale bars: 10µm, except 50µm for panels A, D. Apical-basal (A-B) is indicated for each close up illustrating an epithelial tissue. In box plots, the black bar indicates the median, the whiskers indicate the maximal range, and each value is indicated by a dot. Statistical tests: T-test in F, G. p-value: *, <0.05; **, <0.01; ns, non-significant).

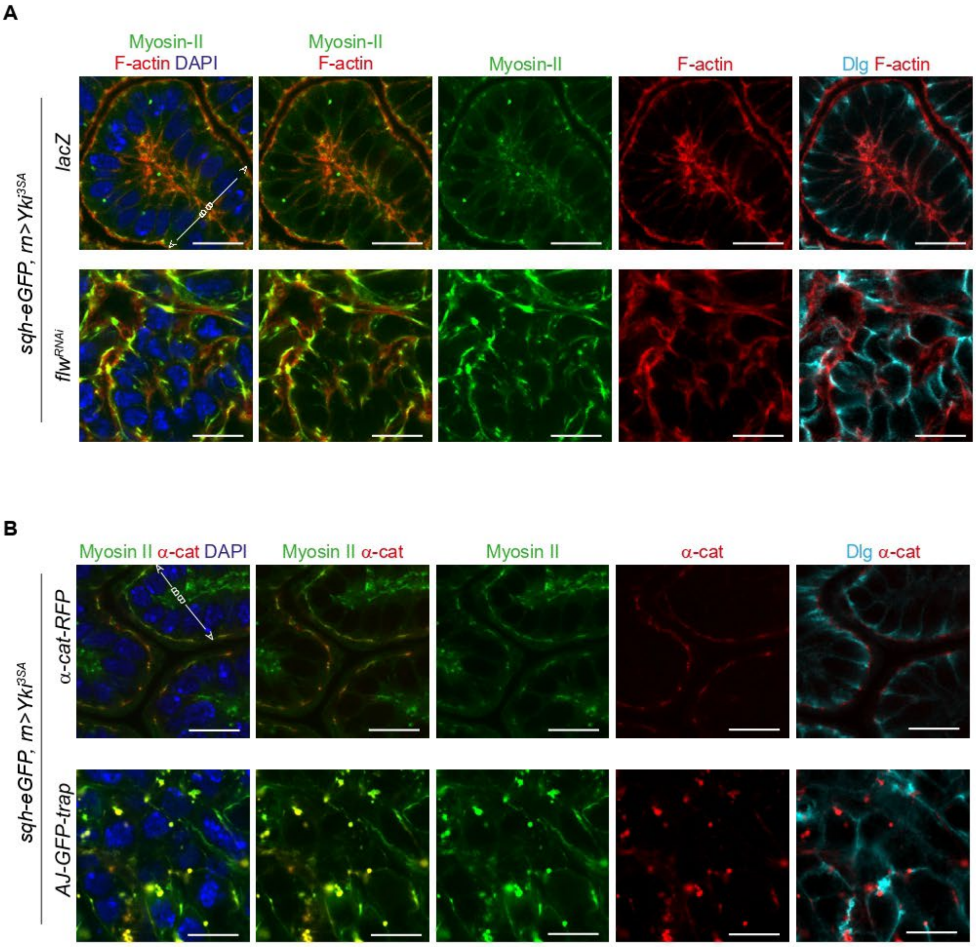


**Supplementary Fig. 4**

**Supplementary Figure 4. Hyperplasia-to-dysplasia transition caused by increased adherens junction tension in Yorkie tumours**

(A) Close up views of the organisation of Yorkie tumours either control (lacZ, top) or in which myosin II levels were increased due to flapwing RNAi expression (bottom). Myosin II (green) levels are increased in mesenchymal cells (bottom), compared to control epithelial cells (top), and reorganize discontinuously around the cell cortex. Nuclei are labelled in blue (DAPI staining), F-actin in red and tight junctions in cyan (Dlg). Number of discs analysed: lacZ, 3; flapwing RNAi, 6. (B) Close up views of the organisation of Yorkie tumours without (alpha-Catenin-RFP expression, top) or with myosin II increase at adherens junction due to the expression of the AJ-GFP trap (bottom). Nuclei are labelled in blue (DAPI staining), adherens junctions in red (alpha-Catenin-RFP), tight junctions in cyan (Dlg) and endogenous myosin II-GFP in green. Number of discs analysed: alpha-CatRFP, 30; AJ-GFP-Trap, 17.

Transgenes are driven in the wing pouch using the *rotund*::Gal4 driver at 25°C for flapwing RNAi (B) or 22°C for AJ-GFP trap experiments. Scale bars: 10μm. Apical-basal (A-B) is indicated for each close up illustrating an epithelial tissue.

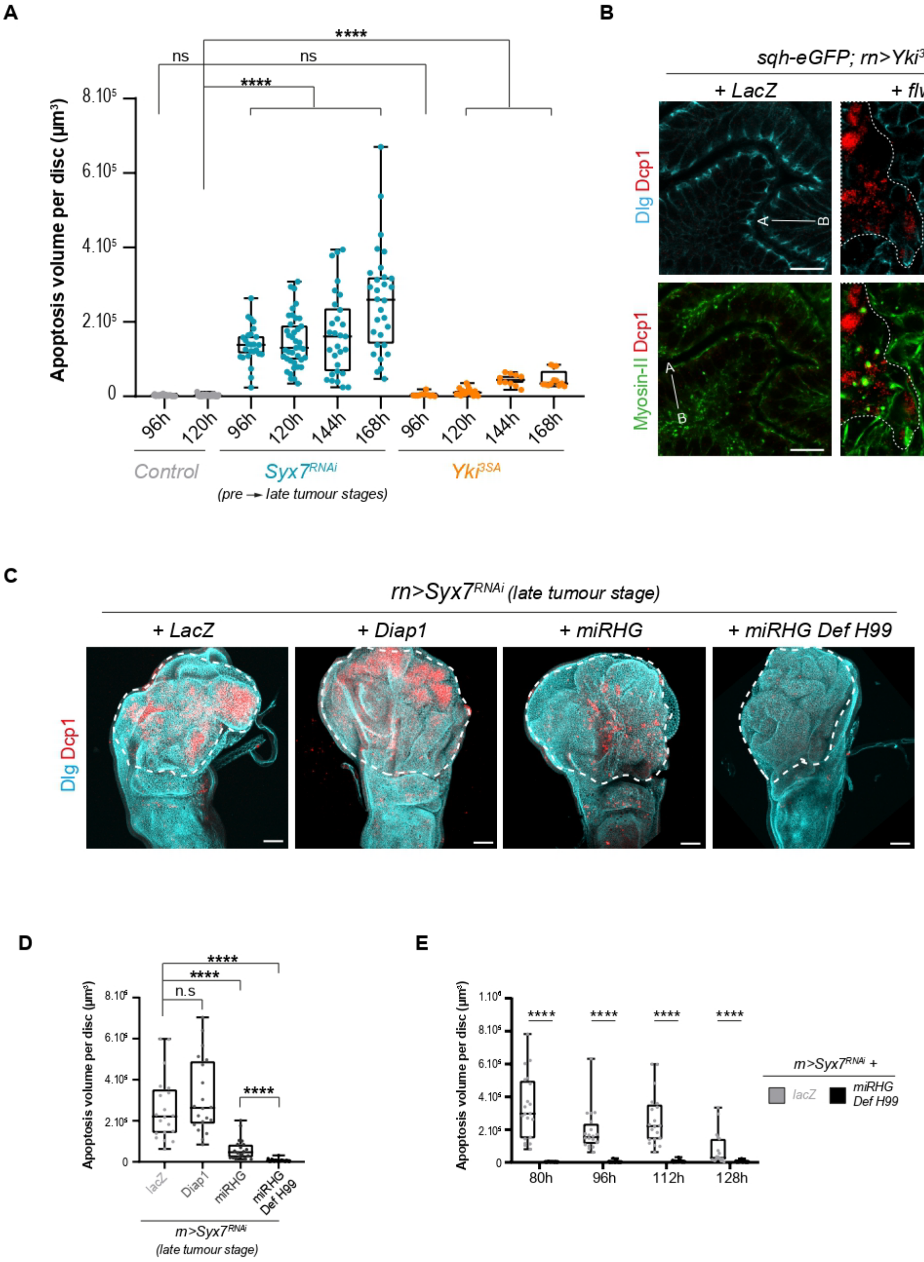


**Supplementary Fig. 5**

**Supplementary Figure 5. Characterisation and manipulation of apoptosis in fly tumours**

(A) Quantification of apoptosis in wing discs in control (grey), Syx7 RNAi (blue) and Yorkie overexpression (orange) conditions at the indicated timepoints (+/- 1.5h AEL). Number of discs analysed: 26 and 38 in control discs at 96h and 120h AEL respectively; 26, 39, 29 and 30 at 96h, 120h, 144h and 168h AEL respectively in Syx7 RNAi, and 11, 18, 10 and 12 at 96h, 120h, 144h and 168h AEL respectively in Yorkie tumours. Experiments were conducted at 25°C. (B) Close up views showing myosin II (green), Dlg (cyan) and active caspase DCP-1 (red) in Yorkie tumours either expressing lacZ (control, left, n=14) or *flapwing* RNAi to increase myosin II levels (right, n=16). Note that mesenchymal cells were in close vicinity of apoptotic domain following myosin II increase. (C) General views (projections) of Syx7 RNAi tumours coexpressing lacZ (control, n=20), Diap1 (n=20), miRHG alone (n=20) or miRHG in a heterozygous background for the H99 deficiency (miRHG Def H99, n=20) at the late stage (112h AEL at 30°C). Tumour shape is visualized using Dlg (cyan), while apoptosis is revealed using anti-active Dcp-1 caspase staining (red). Syx7 RNAi tumours have a similar size and shape, irrespective of apoptosis inhibition efficiency. The pouch domain is indicated by a dashed line. Scale bar: 50µm. (D) Quantification of apoptosis at the late tumour stage (112h AEL at 30°C) for the indicated genotypes. Apoptosis is not affected by Diap1 overexpression, reduced but not abolished by miRHG expression, and very low in miRHG Def H99. (E) Quantification of apoptosis in Syx7 RNAi tumours with (lacZ, in grey) or without apoptosis (miRHG Def H99, black) at the indicated timepoints at 30°C. Note that apoptosis is always efficiently abolished in Syx7 RNAi miRHG Def H99 tumours, irrespective of the timepoint analysed. Number of discs analysed are respectively 20, 20, 20 and 19 at 80h, 96h, 112h and 128h AEL in controls and 14, 20, 20 and 22 in miRHG Def H99.

In box plots, the black bar indicates the median, the whiskers indicate the maximal range, and each value is indicated by a dot. Statistical tests: Mann-Whitney in A, D, E. p-value: *, <0.05; **, <0.01; ***, <0.001; ****, <0.0001; ns, non-significant). All statistical tests were performed by comparison with the control at 120h AEL in (A), and between the control and miRHG Def H99 at each indicated timepoint in (E).

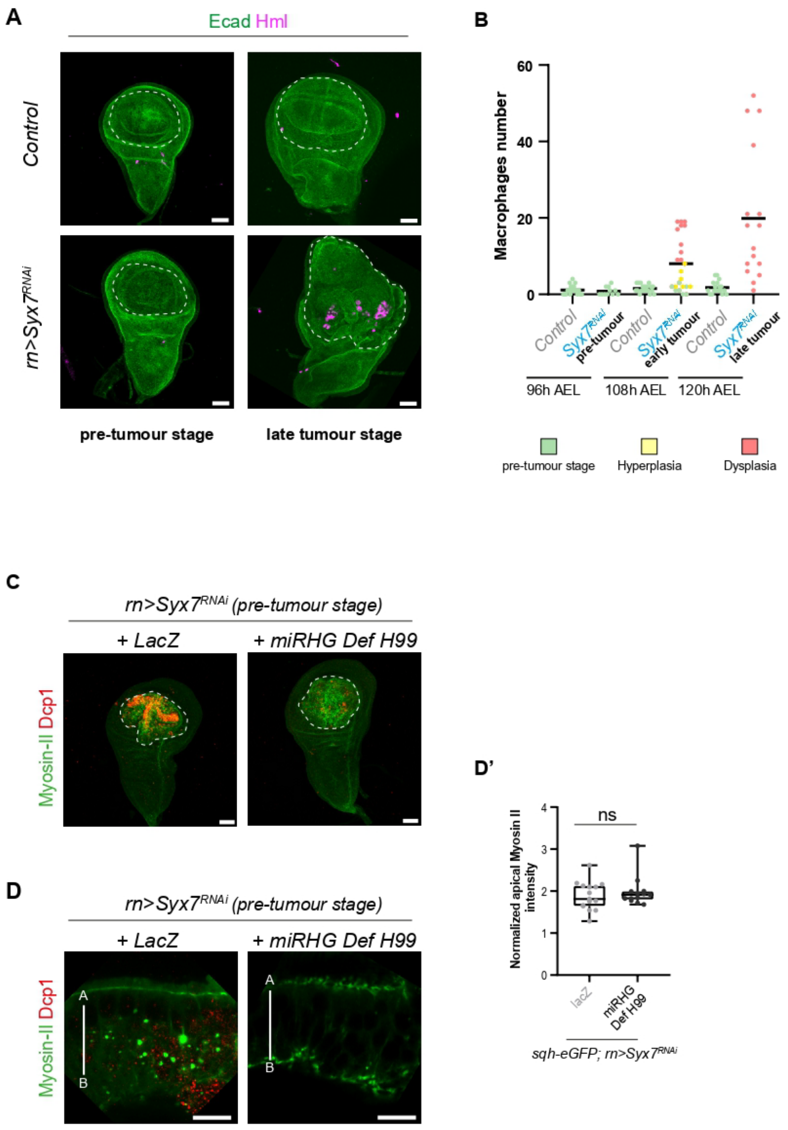


**Supplementary Fig. 6**

**Supplementary Figure 6. Apoptosis does not promote dysplasia formation through macrophage recruitment or increase in tissue tension**

(A, B) Analysis of macrophages (hemocytes) recruitment in Syx7 RNAi tumours. In (A), top and bottom rows show projection views of control and Syx7 RNAi discs respectively. Syx7 RNAi discs are at the pre-tumour stage (96h AEL, left) or late tumour stage (120h AEL, right). ECad (green) labels the discs. Macrophages, labelled with Hml::QF2, QUAS::mCherry transgenes, are shown in magenta. (B) Quantifications of the number of macrophages in the contexts shown in (A). Each dot represents the number of macrophages in one individual disc, and is colour-coded depending on the stage of the disc determined based on ECad staining (size of the wing pouch and epithelial vs mesenchymal state of the cells). Note that macrophages are recruited once tumours have turned dysplastic (red dots at 108 and 120h), but not before the hyperplastic-to-dysplastic switch (green and yellow dots at 96 and 108h). Number of discs analysed in control and Syx7 RNAi discs respectively: 20 and 11 at 96h, 19 and 23 at 108h and 18 and 16 at 120h. (C, D) General views (projections, C) and close up views (single plane, D) of Syx7 RNAi tumours with (lacZ, left) or without apoptosis (miRHG Def H99, right) at a similar age which corresponds to the pre-tumour stage in controls (80h AEL at 30°C). Myosin II (green) accumulates similarly in presence or absence of apoptosis (red) in early Syx7 RNAi discs, apart from apoptotic fragments (bright dots in D). (D') Quantification of myosin II levels at adherens junctions in Syx7 RNAi tumours with (lacZ, grey, n=15) or without (miRHG H99, black, n=13) apoptosis.

Transgenes are driven in the wing pouch using the *rotund*::Gal4 driver at °C in (A, B), or at 30°C in (C-D').The pouch is indicated by a dashed line in C. Apical-basal (A-B) is indicated for each close up illustrating an epithelial tissue. Scale bars: 10µm in D and 50µm in A and C. In box plots, the black bar indicates the median, the whiskers indicate the maximal range, and each value is indicated by a dot. Statistical tests: Mann-Whitney in D'. ns, non-significant.
